# *In vitro* metabolic interaction network of a rationally designed nasal microbiota community

**DOI:** 10.1101/2024.10.23.619785

**Authors:** L. Bonillo-Lopez, O. Rouam-el Khatab, P. Obregon-Gutierrez, I. Florez-Sarasa, F. Correa-Fiz, M. Sibila, V. Aragon, K. Kochanowski

## Abstract

Mounting evidence suggests that metabolite exchange between microbiota members is a key driver of microbiota composition. However, we still know little about the metabolic interaction networks that occur within many microbiota. This is particularly true for the nasal microbiota, and current efforts towards this end are hampered by a lack of microbial consortia that would enable the mapping of metabolic interactions between nasal microbiota members under *in vitro* conditions. To tackle these issues, we developed the Porcine Nasal Consortium (PNC8), a rationally designed microbial consortium of eight strains representing the most *in vivo* abundant genera in the nasal microbiota of healthy piglets. We used this consortium to systematically examine the metabolic capabilities of nasal microbiota members, as well as the metabolic interactions occurring between them. We found that PNC8 strains differ substantially in their metabolic pathway repertoire and ability to grow across various *in vitro* conditions. Nevertheless, spent-media experiments revealed that most metabolic interactions between PNC8 strains are negative, and exometabolomics data pointed to co-depletion of sugars as a key driver of this interaction network. Finally, direct co-cultivation experiments showed that, as a result of this largely negative metabolic interaction network, competition is common among pairs of PNC8 strains and leads to a complex competition hierarchy in which only few strains are able to consistently outcompete all others. Overall, this work provides a valuable resource for studying the nasal microbiota under experimentally tractable *in vitro* conditions and is a key step towards mapping its metabolic interaction network.

## Introduction

Animals are hosts to a plethora of microorganisms occupying virtually all accessible body sites, collectively known as the commensal microbiota ^1^. Given the importance of the commensal microbiota in maintaining host health, there have been extensive efforts to understand the mechanisms that determine microbiota composition. Recent evidence, largely stemming from studies of the gut microbiota, points to metabolic interactions within the microbiota as a key component ^2–6^. These studies suggest that metabolic interactions within the gut microbiota are pivotal not only for determining its composition, but also for providing protection against pathogen invasion by “blocking” the nutrients available to pathogens ^4^. However, it is unclear whether metabolic interactions play a similarly important role in the microbiota of other body sites. This is particularly true for the nasal microbiota, which is the first hurdle many respiratory bacterial pathogens need to overcome when colonizing the host ^7^. Currently, we know little about the metabolic interactions that occur within the nasal microbiota ^8^, and efforts to map such interactions are hampered by a lack of adequate experimental tools, and in particular the lack of synthetic consortia. Synthetic microbial consortia, such as the Oligo-Mouse Microbiota (OMM12) ^9^, have been critical in enabling the identification of metabolic interactions within the gut microbiota ^10–16^. However, the few currently available nasal communities ^17,18^ only cover a small fraction of the diversity present in the *in vivo* nasal microbiota.

To address these issues, we developed the Porcine Nasal Consortium (PNC8), a defined microbial consortium that consists of eight strains representing the most *in vivo* abundant genera in the nasal microbiota of healthy piglets. Here, we used this consortium to systematically characterize the metabolic capabilities of nasal microbiota members and generate the first *in vitro* map of their metabolic interactions, using a combination of exometabolism, spent-media experiments, and direct co-cultivation. Despite substantial differences in metabolic capabilities between PNC8 members, most of their interactions were negative, and frequently driven by competition for metabolites (in particular sugars). This prevalent metabolite competition between PNC8 strains resulted in a highly competitive interaction network with a complex competition hierarchy, in which only few strains were able to outcompete the others. Thus, this work not only provides a valuable new resource for studying the nasal microbiota under *in vitro* conditions, but also presents the first systematic map of its metabolic interaction network.

## Results

### Section 1: developing a defined consortium of the pig nasal microbiota (PNC8)

To enable the elucidation of metabolic interactions within the nasal microbiota under experimentally tractable *in vitro* conditions, we aimed to develop a defined consortium of nasal microbiota members. Specifically, we focused on the pig nasal microbiota, motivated by the importance of respiratory bacterial pathogens in pig production, and the large similarities in nasal microbiome composition between pigs and humans ^19,20^.

As a starting point, we identified eight genera that are abundant and prevalent *in vivo* in the nasal microbiota of healthy piglets (**Figure 1A-B**, **see supplementary text 1 for a detailed description of how this consortium was developed**). Collectively, these genera constitute the majority (and up to 90% and more) of the *in vivo* composition of the pig nasal microbiota as detected in different published data sets (**Figure 1C and Supplementary Figure 1**). We selected a single bacterial strain to represent each of these taxa, as a trade-off between recapitulating the nasal microbiota complexity found *in vivo* and ensuring experimental tractability (following design principles laid out in the development of other defined consortia, e.g. ^9,21^). Wherever available, we used previously characterized ^22–25^ strains (e.g. *M. pluranimalium* LG6-2 ^23^ and *G. parasuis* F9 ^25^). For the remaining genera, we used strains previously isolated from healthy animals and selected the representative strain by prioritizing strains that grew *in vitro* and had low pathogenicity traits (such as hemolysis, see full strain list in **Supplementary Table 1**). The result of these efforts is the **Porcine Nasal Consortium (PNC8)** consisting of eight genetically diverse strains (**Figure 1D**) that collectively cover a large fraction of the predicted metabolic repertoire found in the *in vivo* pig nasal microbiota (as determined before ^22^) (**Figure 1E**). Thus, these analyses suggested that this consortium is a good representation not only of the overall genetic diversity (in terms of included taxa), but also of the metabolic capabilities present in the pig nasal microbiota.

**Figure 1.**
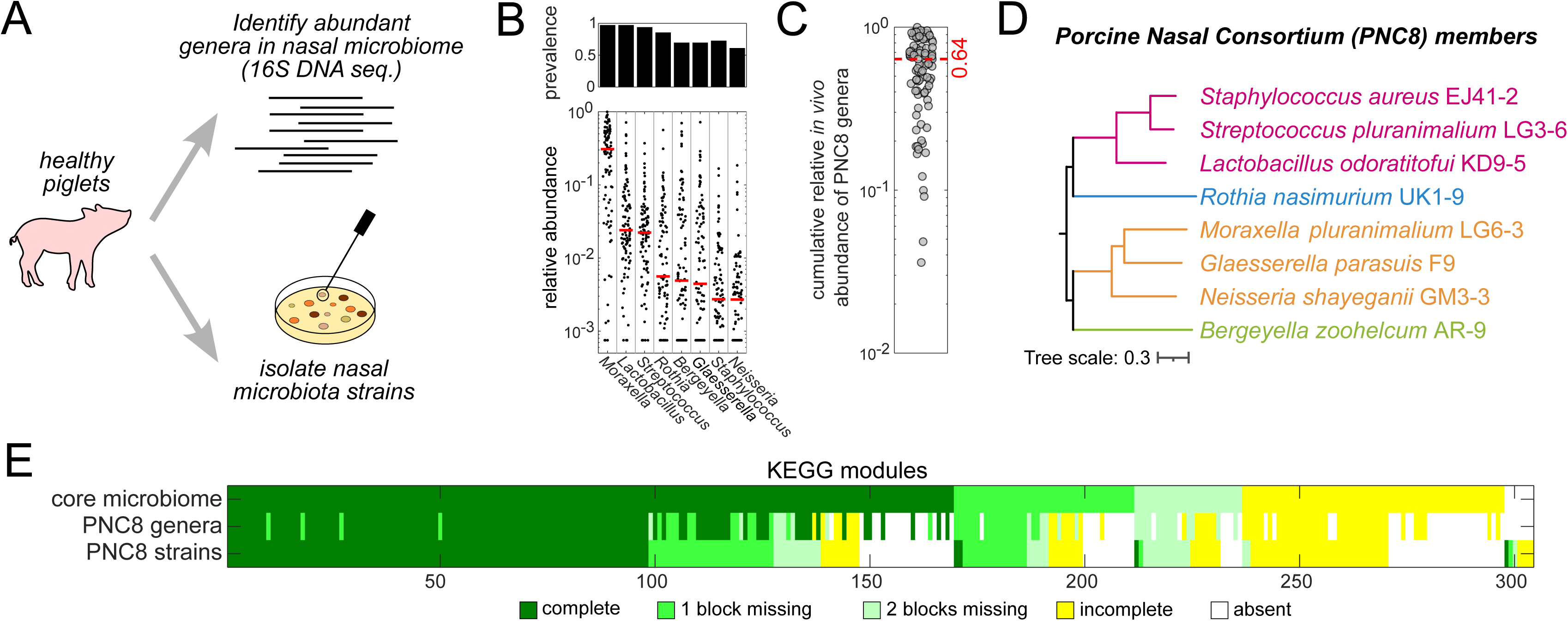
Development of the Porcine Nasal Consortium (PNC8). **A)** Schematic of approach to rationally design the PNC8 consortium. **B)** Prevalence (fraction of samples with relative abundance > 0.1%, top) and relative abundance (bottom) of the eight most abundant genera (sorted by median abundance across samples, shown as red horizontal lines) in 94 healthy piglets as determined by 16S sequencing. Each circle denotes an individual animal. **C)** Summed *in vivo* relative abundance of PNC8 genera (dashed line denotes median: 64%). Each circle denotes an individual animal. **D)** Phylogenetic tree of PNC8 members. PNC8 members are color-coded by phylum (Purple: *Firmicutes*, Green: *Bacteroidota*, Blue: *Actinobacteria*, Orange: *Proteobacteria*). **E)** Comparison of inferred KEGG completeness within *in vivo* samples (inferred using core nasal microbiota or only PNC8 genera based on 16S ASVs, top and middle rows, see methods), and the combined PNC8 strains (see A, bottom row). To aid visual comparison KEGG modules were sorted according to completeness first for core nasal microbiota, and then for PNC8 strains.

### Section 2: characterizing in vitro growth capabilities of individual PNC8 members

Next, we aimed to characterize the metabolic capabilities of the PNC8 strains in more detail. Genomic analysis revealed that these strains vary substantially in their repertoire of metabolic pathways (between 21 and 52 complete KEGG metabolic modules present, see **Supplementary Figure 2**). To test whether these differences in metabolic pathway presence across strains were also reflected in different *in vitro* growth patterns, we examined the ability of each strain to grow in 23 metabolic diverse *in vitro* cultivation media of varying composition and richness (see full list of conditions in **Supplementary Table 2**). These experiments confirmed previously known metabolic requirements of the respective genera, for example the inability of *Glaesserella* species to grow without NAD^+^ supplementation (**Figure 2 and Supplementary Figure 3**). In addition, these experiments revealed substantial differences in growth patterns across PNC8 strains. Typically, any given strain only grew well in one or a few conditions, which also tended to differ across PNC8 strains. Only one strain (*S. aureus* EJ41-2) was able to grow in most tested conditions. Notably, none of the strains grew in a minimal medium supplemented only with glucose (and inorganic nitrogen/phosphor/sulfate), suggesting that all PNC8 strains require at least some external metabolites from their environment. Thus, these data showed that most nasal microbiota members (as represented here by the PNC8 strains) tend to be metabolic “specialists” with distinct nutrient requirements.

**Figure 2.**
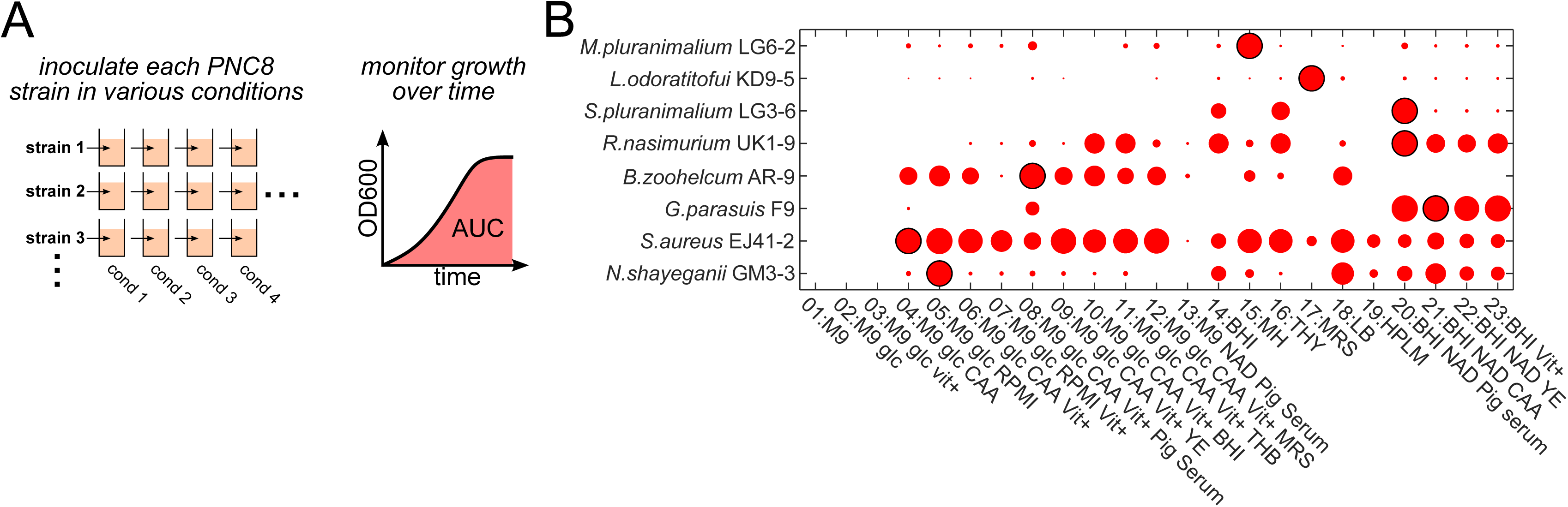
Growth patterns of PNC8 members across various *in vitro* conditions. **A)** Schematic of approach. **B)** Area-under-growth-curve (AUC, shown here as circle size) of PNC8 members in 23 different cultivation media (normalized to the condition with maximal AUC for each PNC8 member, which is denoted with a black edge). See **Supplementary Figure 3** for the underlying growth curves.

### Section 3: examining metabolic activity of PNC8 strains in the same in vitro condition

To elucidate to which extent these distinct nutrient requirements also manifest as distinct metabolic activity in the same condition, we selected a single condition (Brain-Heart Infusion broth supplemented with 80 µg/mL NAD^+^ and 1% inactivated pig serum, termed BHI+ here, condition 20 in **Figure 2**) that supported robust growth in most PNC8 strains. To examine metabolic activity in other nasal microbiota members beyond the PNC8 strains, we also tested several additional commensal strains (belonging to the same PNC8 genera), as well as three pathogenic strains (i.e. *S. suis* P1/7*, G. parasuis* Nagasaki*, A. pleuropneumoniae* 4074). We found that biomass production and pH changes (as general markers of metabolic activity) varied widely across strains (**Supplementary Figure 4A**). Nevertheless, most strains acidified the growth media proportionally to their biomass production (**Supplementary Figure 4B**), suggesting that fermentative metabolism is common among nasal microbiota members in the tested condition. One exception was the PNC8 strain *M. pluranimalium* LG6-2, which did not alter medium pH despite substantial biomass production.

To obtain a more detailed picture of each strain’s metabolic activity, we further quantified concentration changes of about 50 extracellular metabolites (using exometabolomics, see schematic in **Figure 3A** and methods). These data revealed complex metabolite usage patterns that differed substantially across metabolite classes (**Figure 3** and **Supplementary** Figures 5A **& 6**). Moreover, in those cases where we had data available for more than one strain from the same genera included in the PNC8 (i.e. *Rothia*, *Streptococcus*, *Staphylococcus*), the metabolite usage patterns were largely conserved at the species level (or to a lesser extent, at the genus level, see **Supplementary Figure 5B-C**). Many metabolites, and in particular amino acids, remained largely constant in most tested strains (see e.g. glutamine in **Figure 3B**), suggesting that reliance on amino acid catabolism is rare among nasal microbiota members. Some metabolites were strongly depleted in individual strains but remained constant in most others (e.g. AMP in **Figure 3B**), pointing towards specific metabolic requirements. Finally, a subset of metabolites, in particular sugars, were consumed by most strains (e.g. glucose in **Figure 3B**). This point became even clearer when re-sorting the metabolites according to their consumption/production patterns (**Figure 3C**). Nevertheless, none of these metabolites were consumed by **all** tested strains: For example, glucose, which is the main sugar in this cultivation media and was depleted by most strains, was not consumed by the PNC8 strain *M. pluranimalium* LG6-2. This finding is consistent with the observed lack of media acidification by this strain (see **Supplementary Figure 4**) and suggests that there is no single “universal metabolic currency” within the nasal microbiota. Moreover, only few metabolites were both secreted and consumed (the most notable exception was succinic acid, see **Figure 3B**), indicating limited cross-feeding potential.

**Figure 3.**
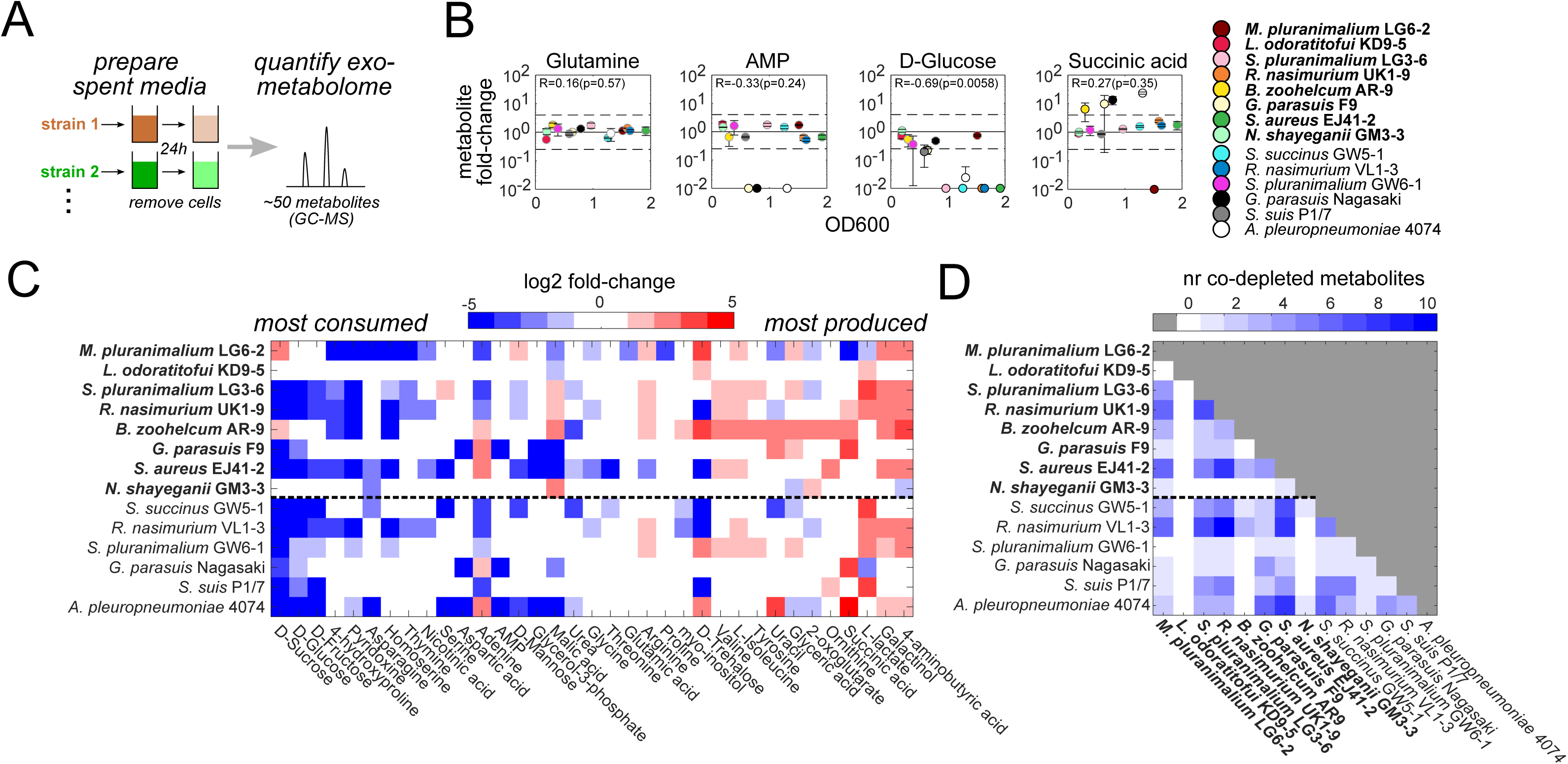
Metabolite usage patterns of nasal microbiota members. **A)** Schematic of experimental design. **B)** Example data, shown here as metabolite fold-changes (compared to fresh media) plotted against the mean OD600 reached in the respective spent-media culture. Error bars denote standard deviation (n = 2). Horizontal dashed lines denote a four-fold change (used to determine depleted/produced metabolites in **C** and **D**). R and p denote the Spearman correlation coefficient and corresponding p-value between metabolite fold-changes and OD600. Data points with a metabolite fold-change below 10^-2^ were set to 10^-2^. See **Supplementary Figure 6** for all other metabolites. **C)** Metabolites sorted from most consumed (i.e. used as substrate by most strains) to most produced (used as product by most strains). Data shown are log_2_ fold-change compared to fresh media (average of two replicate cultures). Unused metabolites (abs(fold-change) < 4 in all tested strains) are not shown. **Bold strain names**: PNC8 strains. Data shown: average of two replicate cultures. **D)** Number of co-depleted metabolites (>4-fold decrease compared to fresh media) for all tested strain pairs. **Bold strain names**: PNC8 strains.

To examine the extent to which nasal microbiota strains overlap in their metabolite requirements, we further calculated the number of co-depleted metabolites in all possible strain pairs (that is, metabolites that were depleted at least four-fold by both strains when grown in isolation). Even within this relatively small set of 50 metabolites examined here, co-depletion was prevalent across strain pairs (**Figure 3D**). A notable exception was the PNC8 strain *Lactobacillus odoratitofui* KD9-5, whose metabolite depletion pattern largely did not overlap with any of the other strains. Overall, these experiments revealed complex metabolite usage patterns in nasal microbiota members, with prevalent (albeit not universal) depletion of sugars. In particular, our data revealed that despite their distinct metabolic pathway repertoire and growth patterns across *in vitro* conditions, many nasal microbiota members do substantially overlap in the metabolites they consume *in vitro*.

### Section 4: identifying directional metabolic interactions between PNC8 members using spent-media experiments

The exometabolomics data described above suggested that most PNC8 strains have the potential to compete at least for some nutrients, as indicated by the prevalence of co-depleted metabolites. However, since many of these strains also secreted at least one metabolite in the tested condition, it is conceivable that these secretion products could serve as alternative nutrients for other strains and thereby alleviate this potential nutrient competition. To examine the potential for competition between PNC8 strains more directly, we therefore used spent-media experiments, which enable the detection of directional metabolic interactions between strain pairs by monitoring their ability to grow in their respective partner strain’s spent media ^10,13,26,27^ (see schematic of experiment in **Figure 4A**). As before, we selected a single condition that supports robust growth of most PNC8 strains (i.e. BHI+). To disentangle the impact of direct (i.e. caused by changes in metabolite concentrations) and indirect (i.e. caused by changes in pH) metabolic effects ^13,27^, we performed these experiments both with and without re-adjusting the pH back to a range between 7.2-7.4 (the pH of fresh BHI+ medium).

**Figure 4.**
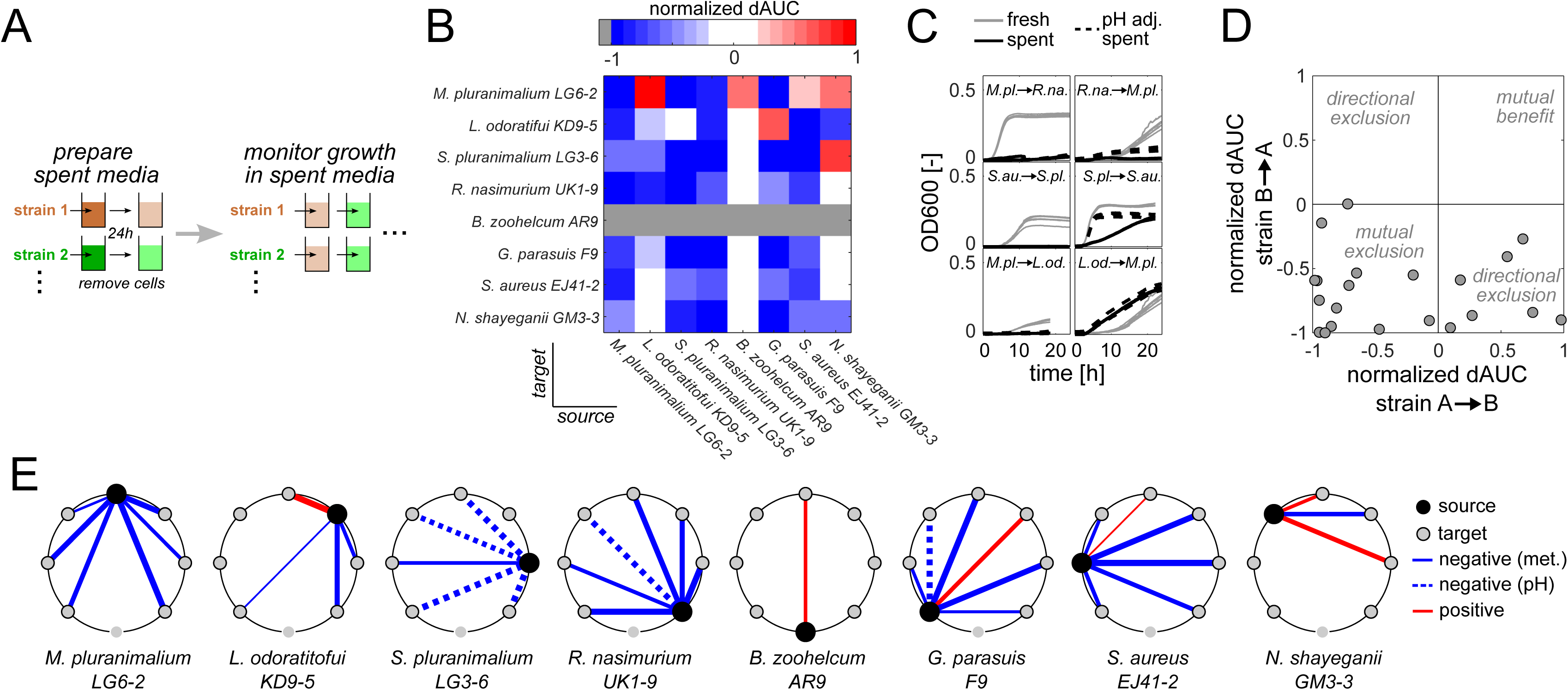
Identifying directional metabolic interactions between PNC8 strains using spent-media growth experiments. **A)** Schematic of experimental approach. **B)** Normalized difference in area-under-the-growth-curves (dAUC). Normalized dAUC = (AUCspent – AUCfresh) / AUCfresh) for all PNC8 strains. Negative normalized dAUC values denote cases where a strain grows more poorly in a spent media compared to fresh media. Data show mean of 2-3 replicate cultures. **C)** Example growth curves (shown from source → target strain). Each curve denotes an individual replicate curve (n = 2-3 for spent media, n = 6 for fresh media). **D)** Same data as in **B**), but plotted to highlight directional impact in each strain pair. **E)** Pairwise interaction maps extracted from growth patterns in spent media, shown separately for each PNC8 strain as a source. Line thickness denotes the interaction strength (weak interactions with dAUC between −0.25 and 0.25 were omitted). Interactions were labeled as pH-driven (dashed lines) if the difference in dAUC between normal and pH-adjusted spent media exceeded 0.55. Note that *B. zoohelcum* AR-9 grew neither in fresh nor in spent media and was therefore omitted from the analysis. See **Supplementary Figure 8** for the underlying growth curves.

Generally, the pairwise interaction network of PNC8 strains was dominated by negative interactions, in which strains typically grew more poorly in most tested spent media (**Figure 4B**, **Supplementary Figure 7** and **Supplementary Figure 8**). In most cases, these interactions were either bidirectionally negative, (e.g. *R. nasimurium* UK1-9 and *M. pluranimalium* LG6-2 in **Figure 4C**), or negative in one direction. However, comparison of pH-adjusted and non-adjusted data suggested that not all of these interactions were purely nutrient-driven: for example, the negative impact of *S. pluranimalium* LG3-6 (which showed the strongest degree of media acidification among the PNC8 strains, see **Supplementary Figure 4A**) on other PNC8 members was largely abolished after re-adjusting the pH of its spent media (**Supplementary Figure 8**, see example in **Figure 4C**). Nor were the interactions between PNC8 members exclusively negative: there were a few instances in which the growth of a strain was improved in the spent media of another (e.g. *M. pluranimalium* LG6-2 in the spent media of *L. odoratitofui* KD9-5, **Figure 4C** and **Supplementary Figure 8**), even though none of these positive interactions were mutual (**Figure 4D**). Overall, these spent-media experiments revealed a complex network of mostly negative interactions between PNC8 strains, which differed both in terms of interaction strength as well as mode of operation (i.e. purely metabolic versus pH-driven, see summary in **Figure 4E**).

### Section 5: examining the impact of metabolite competition on PNC8 co-cultivation outcome using pairwise co-cultivation assays

The spent-media growth experiments revealed that negative (and largely bidirectional) interactions were prevalent between PNC8 strains, and pointed to metabolite competition as the main driver of this interaction network. Next, we wanted to examine the impact of such prevalent metabolite competition on situations where these strains are cultivated together.

To better understand the potential impact of metabolite competition on co-cultivation outcomes, we simulated (pairwise) co-cultivation experiments, in which strains compete for a single growth-limiting metabolite (**Supplementary Text 2**). In line with seminal experimental co-cultivation studies (e.g. ^28^), these simulations predicted that metabolite competition between two strains manifests in a co-culture biomass production that is lower than the summed biomass production of each individual strain when grown in isolation (**Supplementary Figure 9**). To test these predictions experimentally, we performed co-cultivation experiments for all possible PNC8 pairs (28 unique combinations in total) in the same nutrient-rich condition (i.e. BHI+) we had used in the spent-media experiments described above, monitoring the growth of the co-cultures and the respective individual strains over time (schematic of approach shown in **Figure 5A**). When comparing measured and expected biomass production (using the maximal OD600 during 24h co-cultivation as our metric), we found that indeed the vast majority of PNC8 strain pairs had lower than expected co-culture biomass production, suggesting that most pairwise PNC8 strain interactions are competitive in the test condition (**Figure 5B**). Those strain combinations whose measured biomass production did match the expected one mainly included *B. zoohelcum* AR-9, the sole PNC8 strain which does not grow in the tested condition (gray circles in **Figure 5B**).

**Figure 5.**
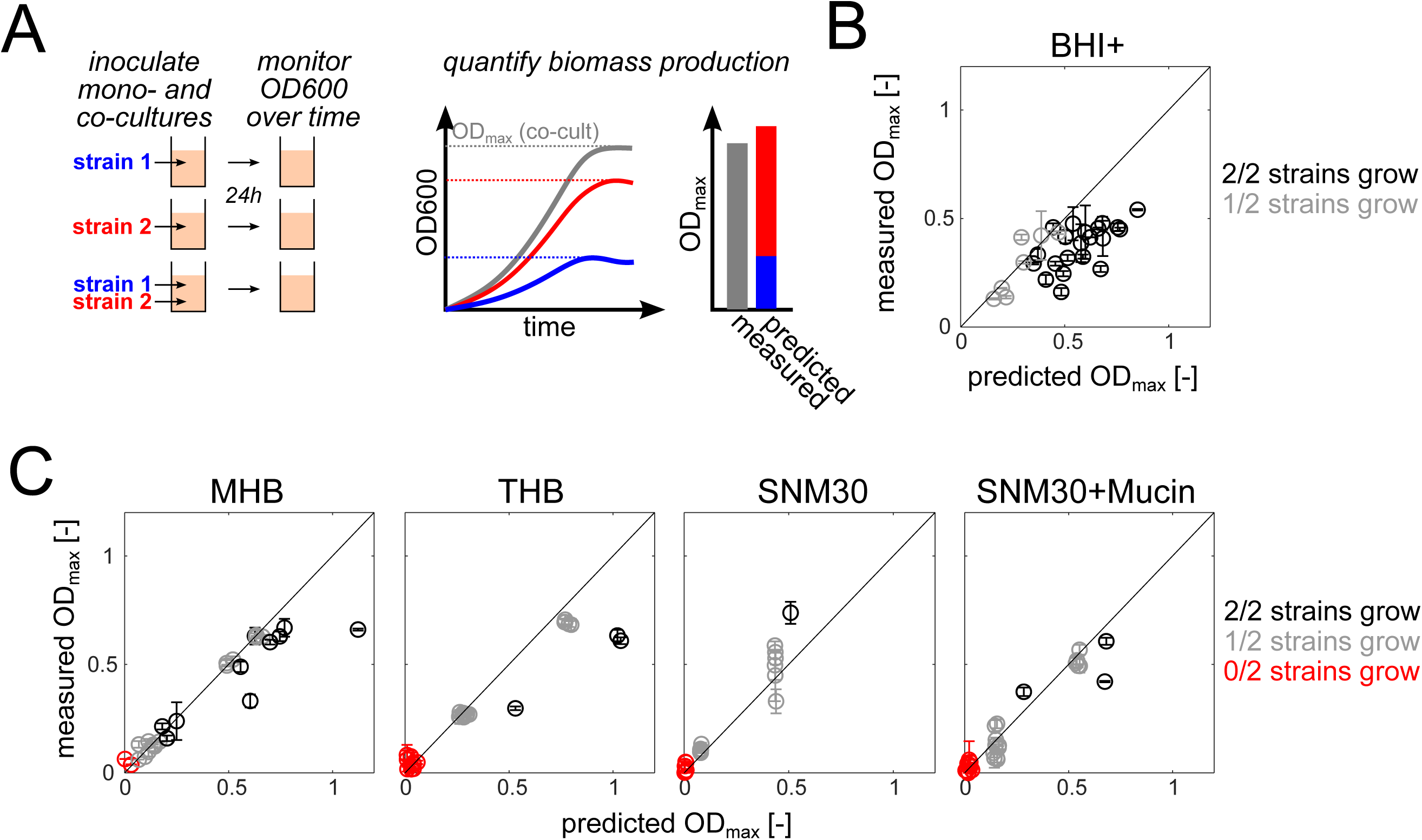
Identifying instances of metabolite competition between PNC8 using direct co-cultivation assays. **A)** Schematic of approach. **B)** Predicted (from individual strains) and measured maximal OD600 of all 28 possible PNC8 co-cultivation pairs in BHI+. **C**) Predicted and measured maximal OD600 of all co-cultivation pairs in four additional conditions. Black: Both strains grow in isolation (defined here as maximal OD600 > 0.1). Gray: only one of the two strains grows in isolation. Red: neither of the two strains grows in isolation. Error bars denote standard deviation (n = 2). See **Supplementary Figure 11** for the underlying growth curves.

Next, we wanted to test whether this prevalence of metabolite competition between PNC8 strains is a unique feature of the tested condition (i.e. BHI+). Towards this end, we repeated these pairwise co-cultivation experiments in four additional media differing in composition and richness, namely Todd-Hewitt-Broth (THB) and Muller-Hinton-Broth (MHB) as representatives of commonly used rich cultivation media, as well as two defined media resembling the metabolic composition of human nasal secretions (i.e. synthetic nasal medium, SNM ^29^, tested here both with and without additional 5 g/L mucin). These additional cultivation media were less growth-permissive than the originally chosen condition, with fewer PNC8 strains being able to grow in isolation. Nevertheless, most cases in which both PNC8 strains were able to grow had lower-than-predicted biomass production (black circles in **Figure 5C**), suggesting again the prevalence of metabolite competition. Moreover, we also did not observe cases of robust growth for any of those combinations in which neither PNC8 strain was unable to grow individually, suggesting the absence of “metabolic niche expansion” ^30^ among PNC8 pairs under the tested conditions. Nevertheless, among those combinations in which only one strain was able to grow individually, we did observe (especially in SNM30) isolated cases in which the co-culture biomass production did exceed the prediction (gray circles in **Figure 5C**). A commonly used alternative biomass production metric (area-under-the-growth-curve, AUC ^31^) yielded similar results (**Supplementary Figure 10**).

Overall, these pairwise co-cultivation experiments suggested that – consistent with the findings from our spent-media experiments – the interaction network between PNC8 strains is largely dominated by metabolite competition. Finally, we aimed to test whether our pairwise co-cultivation experiments could not only identify instances of competition, but also reveal which strain outcompeted the other. Towards this end, we made use of the fact that the PNC8 strains have easily distinguishable growth curves in the tested conditions. Therefore, we reasoned that if one strain strongly outcompetes the other, their resulting co-culture growth curve would closely resemble this strain’s individual growth curve. To test this conjecture, we calculated for each PCN8 strain pair and condition (28 pairs x 5 conditions) the relative distance between co-cultures and individual strain growth curves (see schematic approach in **Figure 6A**). This analysis indeed identified many cases in which the co-culture strongly resembled one of the strain’s monoculture (suggesting that this strain strongly outcompetes the other in the tested condition, **Figure 6B**). For some strain pairs (e.g. *M. pluranimalium* LG6-2:*G.parasuis* F9, see **Supplementary Figure 11**), the co-culture resembled either strain 1 or 2 depending on the condition, suggesting that the PNC8 competition hierarchy (that is, which strain is predicted to outcompete which others) is at least to some degree condition-dependent. One exception was *S. aureus* EJ41-2: virtually all co-cultures including this strain strongly resembled its monoculture, indicating that *S. aureus* EJ41-2 outcompetes all other PNC8 strains across all tested conditions (see **Figure 6C**).

**Figure 6.**
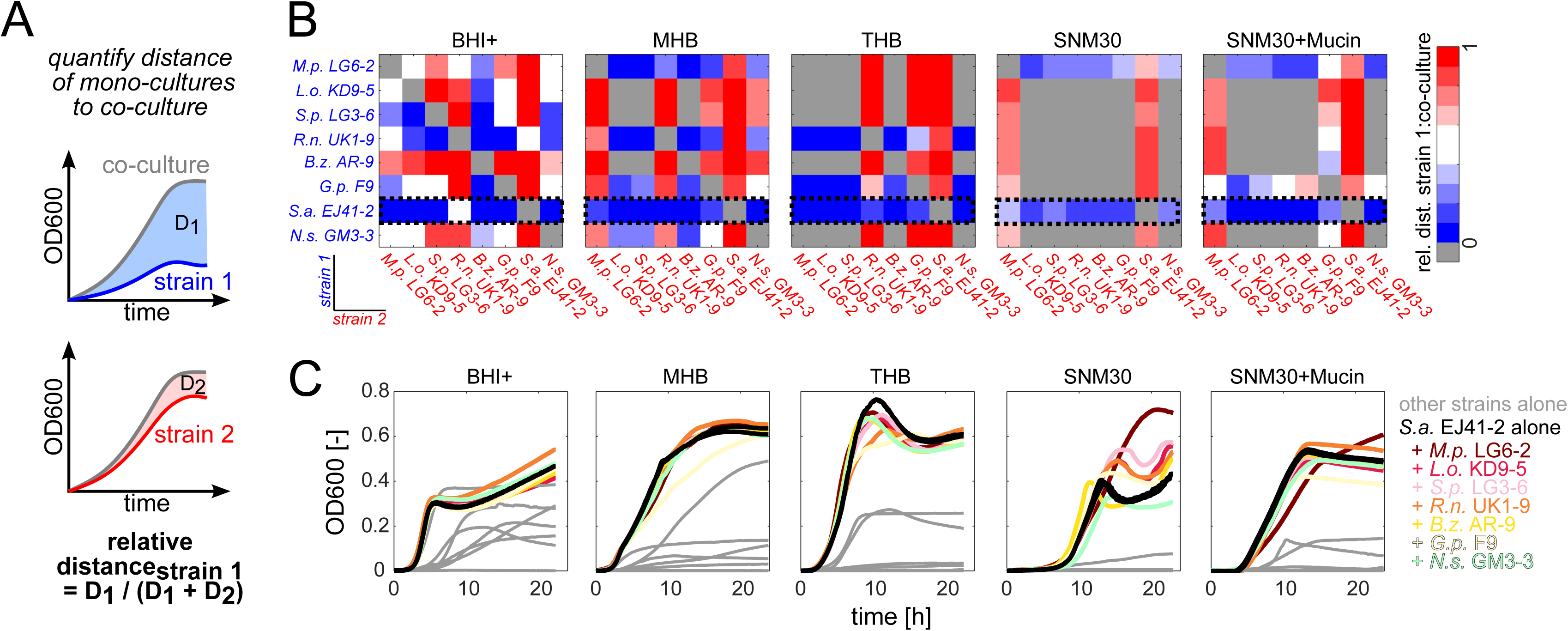
Inferring pairwise competition hierarchy in PNC8 pairwise co-cultures by comparing mono- and co-culture growth curves. **A)** Schematic of analysis approach. **B**) Relative distance of the growth curve of strain 1 to the co-culture growth curve (calculated as described schematically in A) for all strain pairs and conditions. Data shown: mean relative distances across replicates (n = 2). Only pairs in which at least one of the strains grew were considered (the rest is shown in gray). Black boxes with dashed outline: Example strain highlighted in next panel. **C**) Example growth curves for co-cultures including *S.aureus* EJ41-2 across all tested conditions. Black line: growth curve of *S.aureus* EJ41-2 alone. Colored lines: co-culture growth curves of EJ41-2 and other PNC8 strains. Grey lines: growth curves of other PNC8 strains in isolation. Data show mean of two replicate cultures.

What could be a potential explanation for this complex competition hierarchy? The most parsimonious explanation are growth rate differences between the PNC8 strains: our aforementioned simulations of strain pairs competing for growth-limiting metabolites predicted that, in absence of any additional effects, the relative distance of each strain’s individual growth curve to the co-culture curve would be directly proportional to their growth rate differences (that is, the larger the difference in growth rate, the more the co-culture is predicted to resemble the faster strain’s growth curve, see **Supplementary Text 2**). We found that the experimental data largely matched these predictions (**Supplementary Figure 12**): cases in which the co-culture growth curve strongly resembled one of the strains indeed tended to be cases in which that strain grew much faster than the other in isolation (with a few notable exceptions, such as the *R. nasimurium* UK1-9: *M. pluranimalium* LG6-2 pair in BHI+).

Taken together, these systematic pairwise co-cultivation experiments not only revealed that competition is prevalent among PNC8 strains, but also suggested a simple underlying mechanism determining the observed competition hierarchy: For any given strain combination, the ability of one strain to outcompete the other largely depends on their difference in growth rate when grown alone.

## Discussion

In this study, we aimed to elucidate the metabolic interaction network of the nasal microbiota, a microbial community that serves as a critical first hurdle against respiratory pathogen invasion. To enable the study of metabolic interactions within the nasal microbiota under experimentally tractable *in vitro* conditions, we developed a microbial consortium, which we term Porcine Nasal Consortium (**PNC8**). This consortium consists of 8 strains representing the most abundant genera in the nasal microbiota of healthy piglets and covers a large fraction of not only the *in vivo* nasal microbiota composition, but also its metabolic pathway repertoire. Using this consortium, we learned three key lessons about the metabolic interaction network of the nasal microbiota: First, growth assays across various *in vitro* cultivation media indicated that PNC8 strains differ substantially in their metabolic capabilities and nutrient requirements. Second, despite these differences in metabolic capabilities, spent-media experiments revealed that most metabolic interactions between PNC8 strains are negative. Third, direct co-cultivation experiments revealed that due to this prevalence of negative metabolic interactions, competition is common between pairs of PNC8 strains, and results in a complex competition hierarchy where only few strains are able to outcompete all the others.

The picture emerging from the data presented here is that the nasal microbiota is a highly competitive environment, in which most microbe-microbe interactions are negative (see e.g. **Figure 4** and **Figure 5**). This finding is in line with recent studies of gut microbiota communities ^10–15^, and suggests that despite its large differences in composition and lower complexity (e.g. studies of the human nasal microbiota show that only 2-10 species already make up >90% of the nasal microbiota in most individuals ^32^), the nasal microbiota may have similar ecological driving forces. One potential implication is that this prevalence of negative interactions within the nasal microbiota may provide more opportunities for developing microbe-centered strategies against respiratory pathogens, as suggested recently ^33^. In fact, epidemiological studies in humans have shown a negative association between some commensal nasal microbiota members (e.g. *Corynebacterium spp*. and *Dolosigranulum pigrum*) and pathogens such as methicillin-resistant *Staphylococcus aureus* and *Streptococcus pneumoniae* ^34,35^, and seminal work has demonstrated that pathogens could indeed be eliminated from the nasal passage in humans by the introduction of commensal microbiota members ^36^. Future efforts may leverage the metabolic interaction network described here as a starting point for the development of new targeted and metabolically driven intervention strategies against other respiratory pathogens.

Our exometabolomics data also points to some nutrients that may underpin this competitive metabolic interaction landscape. Specifically, our data revealed that sugars such as glucose are being consumed by most nasal microbiota members tested here (See **Figure 3**). This finding is consistent with the fact that many nasal microbiota members (e.g. *Streptococcus* ^37^) rely heavily on fermentative metabolism for energy generation. *In vivo* measurements have shown that there is free glucose (in a high micromolar concentration range) in the human nasal environment ^29^, suggesting that competition for sugars may also be a key driver of metabolic interactions between nasal microbiota members *in vivo*. Interestingly, we found that one PNC8 member, namely *Moraxella pluranimalium* LG6-2, does not actually consume glucose (but rather uses short-chain fatty acids such as succinate as carbon sources). Follow-up experiments suggested that the absence of glucose consumption is a general feature of *Moraxella* strains isolated from the piglet nasal microbiota (**Supplementary Figure S13**). It is tempting to speculate that this ability to “side-step” the fierce competition for sugars may provide *Moraxella* (which is by far the most abundant genus in the *in vivo* piglet nasal microbiota, see **Figure 1B**) with a competitive advantage that enables it to dominate the piglet nasal cavity. Future efforts may examine to which extent the prevalent competition for sugars is also a key feature of the nasal microbiota of other animal species.

Here, we not only show that competition between nasal microbiota members is common, but also that there is a competition hierarchy among strains (where some strains are more competitive than others). Interestingly, the most *in vitro* competitive PNC8 strains, *S. aureus* EJ41-2 and to a lesser extent *R. nasimurium* UK1-9, both belonging to genera that typically show comparatively low abundance in the nasal cavity of piglets (see **Figure 1B**). One potential explanation for this discrepancy is the *in vitro* cultivation media we used here. In fact, the pairwise co-cultivation experiments we performed across different conditions already suggest that the competition hierarchy among nasal microbiota members is at least to some extent condition-dependent (see **Figure 6**). Nevertheless, in all tested conditions *S. aureus* EJ41-2 emerged as the most competitive PNC8 strain (despite the low abundance of this genus in the nasal cavity of piglets). One possible explanation for this discrepancy could be that *in vivo* the abundance of *S. aureus* is being controlled not only through metabolic interactions with other nasal microbiota members, but also by the host itself, for example through the immune system. Future efforts may examine the relationship between the different PNC8 members and the host in more detail, for example by making use of *in vitro* co-cultivation systems that include bacteria as well as porcine nasal organoids ^38^.

This study has several limitations. First, to enable the systematic characterization of nasal microbiota members and their metabolic interactions, we developed a new synthetic consortium (**PNC8**). We did try to ensure that this consortium covers not only the most *in vivo* abundant taxa, but also a large fraction of the metabolic pathways present within the piglet nasal microbiota (see **Figure 1C**). Moreover, our exometabolomics data revealed that at least for those cases we were able to test, other nasal microbiota strains from the same species/genus that were not part of this consortium tended to show similar metabolic activity patterns (see Supplementary **Figure 5B-C**), suggesting that the strains selected here are at least broadly representative of the metabolic capabilities present in the respective genera. Nevertheless, we acknowledge that the consortium we developed here (as any synthetic microbial consortium) cannot capture the full complexity of the nasal microbiota. Future efforts may use complementary approaches, such as the *in vitro* cultivation of nasal microbial communities directly from swabs (as recently demonstrated for the gut microbiota ^39^) to corroborate the findings obtained with our synthetic consortium.

Second, to enable the systematic characterization of the metabolic interaction network within the nasal microbiota, we largely relied on cultivation media that likely do not recapitulate the metabolic environment found *in vivo*. In particular, to be able to systematically examine the metabolic activity of individual PNC8 strains, and the metabolic interactions that occur between them, we needed to use a rich cultivation media (i.e. BHI+) that supports the growth of most PNC8 strains. We did corroborate our findings also in *in vitro* conditions that more closely resemble the nasal metabolic environment found in humans (i.e. the SNM30 and SNM30+mucin conditions used in our pairwise cocultivation experiments, see **Figure 5** and **Figure 6**), and found that only few PNC8 strains were able to growth in these conditions either in isolation or in pairs. Although it is well possible that the nasal metabolic environment of piglets may differ from the human one (and thereby provide some additional metabolites that support the growth of PNC8 strains), we have so far been unable to sample porcine nasal secretions using the experimental procedures developed to sample human nasal secretions (e.g. ^29,40^). Future efforts will be needed to adapt currently available protocols and determine which metabolites are present in the porcine nasal environment.

Finally, in this study we focused on examining the pairwise interaction network of the piglet nasal microbiota (as represented by the **PNC8**). Therefore, we are not able to identify higher-order interactions ^41,42^ between nasal microbiota members (although the importance of such higher-order interactions for microbial communities is a matter of ongoing debate, see e.g. ^43,44^). Future efforts may use our work as a starting point to examine the higher-order interaction space of the nasal microbiota in more detail, for example by making use of new experimental protocols that enable the full factorial construction of multi-member synthetic microbial communities ^45^.

In conclusion, in this work we developed the Porcine Nasal Consortium (PNC8), a new resource for studying the nasal microbiota under experimentally tractable *in vitro* conditions. This consortium enabled us to, for the first time, systematically map the *in vitro* interaction network of the nasal microbiota, and our data suggest that this interaction network is largely negative and dominated by metabolite competition. These findings provide a key step towards understanding the driving forces that shape the nasal microbiota composition.

## Methods

### Reagents and strains

Unless stated otherwise, all reagents were purchased from Sigma-Aldrich. *Moraxella pluranimalium* LG6-2 was originally described in ^23^; *Bergeyella zoohelcum* AR-9 in ^24^, and *Glaesserella parasuis* F9 in ^25,46^. Pathogens included *G. parasuis* Nagasaki (serovar 5 reference strain) originally isolated in Japan from a pig with meningitis ^47^, *Streptococcus suis* P1/7 (serovar 2, gift from Dr. Marcelo Gottschalk); *Actinobacillus pleuropneumoniae 4074* (gift from Dr. Marcelo Gottschalk). All other strains were obtained from our internal strain collection and had original been isolated from chocolate PolyViteX agar plates (Biomerieux) inoculated directly with nasal swabs from 3-4 week old healthy piglets as described before ^23–25,46^. The full list of bacterial strains used in this study is available in supplementary table 1.

### Identification of most abundant nasal microbiota genera

To identify the most abundant genera in the piglet nasal microbiota, 16S sequencing data from 94 nasal swab samples (all taken from healthy farm piglets) from several previous studies conducted in our laboratory and analyzed jointly in a recent publication from our laboratory ^48^ were collated, and taxa were sorted (at genus level) by mean relative abundance across all samples. Gut-microbiota associated taxa (i.e. taxa belonging to *Clostridiales* and *Bacteroidales* orders, see ^48^) were excluded, and the relative abundance of the remaining taxa was recalculated accordingly.

### PNC8 genome sequencing, assembly, and annotation

Genomic DNA was extracted from each PNC8 member strain except *G.parasuis* F9, which had previously been sequenced (NCBI RefSeq Assembly GCF_000731865.1), using a DNA extraction kit (ZymoBIOMICS DNA/RNA Miniprep Kit, Cat# R2002) following manufacturers’ instructions. Three strains (*M.pluranimalium* LG6-2, *R.nasimurium* UK1-9 and *S.pluranimalium* LG3-6) were sequenced with Illumina (short-read sequencing, Novogene) and Nanopore (long-read, Nano1health), while the others (i.e. *N.shayeganii* GM3-3, *S.aureus* EJ41-2, *B.zoohelcum* AR-9 and *L.odoratitofui* KD9-5) were sequenced only with the latter technology. For short-read sequencing, samples were sequenced with Illumina HiSeq, and libraries were prepared using Nextetera XT DNA preparation kit (cat. no. FC-131-1024; Illumina, Inc., San Diego, CA), followed by multiplexed paired-end sequencing (read length of 150 bp). Long Nanopore reads were obtained using a r9.4.1 flowcell (FLO-MIN106) with a Rapid Barcoding Kit (SKQ-RBK004). Basecalling was performed with Guppy version 6.3.8 using its super-accuracy mode (https://nanoporetech.com/document/Guppy-protocol). Since the strains were sequenced at different times, these versions differed. For strains LG6-2 and UK1-9, sequencing was performed in a minION mK1c and Guppy 5.0.16 was used for basecalling. For strain LG3-6, a minION MK1b and the same Guppy version above were used. For strains GM3-3, AR-9, KD9-5 and EJ41-2, a minion Mk1c and Guppy v. 6.2.1 were used. Reads with a score greater than 60 and a length over 200bp were retained. Only high-quality reads from both sequencing platforms were further processed after removing reads with Q-score < 10 using Filtlong 0.2.1 (https://github.com/rrwick/Filtlong). Genome assembly was performed *de novo* with Unicycler version 0.4.8 ^49^ (using hybrid assembly in cases where short- and long-read sequences were available). Assembled genomes were annotated using the bacterial genetic code with RAST tool kit (RASTtk) ^50^ available through the Bacterial and Viral Bioinformatics Resource Center (BV-BRC) ^51^. Subsequently, KEGG functions were inferred for each annotated gene using the eggNOG-mapper ^52^, using translated fasta files as input. In addition, a phylogenetic tree was built from the genomes of all PNC8 members using the “Bacterial Genome Tree” method also available at BV-BRC ^51^ under the default parameters, which uses 100 rounds of the “Rapid” bootstrapping option of RAxML ^53^. A maximum number of 100 genes were selected to be picked randomly, build an alignment, and generate a tree based on the differences within those selected sequences. The resulting phylogenetic tree was visualized and customized using ITOL ^54^ (Version v6.8), where the tree was rooted at midpoint.

### Analysis of KEGG module completeness

KEGG module completeness of individual PNC8 strains was determined with the “reconstruct” function of the KEGG-mapper tool (available online at https://www.genome.jp/kegg/mapper/), using the list of KO identifiers inferred with the eggNOG-mapper (see above), and considering also incomplete modules and modules with missing blocks. To determine KEGG module completeness when considering all PNC8 strains together, the lists of KO identifiers from each strain were first merged before using the KEGG-mapper tool. To infer KEGG module completeness found in *in vivo* samples, a recently identified pig nasal core-microbiota was used as a starting point ^22^, and Amplicon Sequence Variants (ASVs) found in this pig nasal core-microbiota (or only ASVs belonging to the 8 genera included in the PNC8 consortium, see Figure 1) were used to predict KO identifiers using PICRUSt2 ^55–60^ (using its default parameters) and the Kyoto Encyclopedia of Genes and Genomes (KEGG) reference database ^61^. Finally, the resulting KO identifier list was used to determine KEGG module completeness as described above. In each case, results from the KEGG-mapper tool were parsed using a python script available at (https://github.com/JulieMarieCharmillon/kegg_parser) as well as custom bash scripts.

### *In vitro* cultivation experiments

For all *in vitro* cultivation experiments, strains were first plated out on chocolate agar (except *L. odoratitofui*, which was plated on MRS agar) directly from −80°C stocks and were incubated overnight at 37°C in a 5% CO_2_-enriched atmosphere.

#### Growth experiments across metabolic environments

To characterize the growth of bacterial strains across diverse metabolic environments, cell suspensions were freshly prepared from agar plates (at OD600 of 0.3 in sterile PBS) and used 1:30 diluted to inoculate 150 microL cultures of 23 different growth media (listed in **supplementary table 2**) in 96-well plate format (Greiner Cat. No 655 180). Subsequently, 100 *µ*L mineral oil (M3516, Sigma-Aldrich) was added to each well to prevent evaporation ^62^. Bacterial growth (as determined by OD600) was monitored every 10 min for around 24h with an automated plate reader (Tecan Nano M+) using an incubation temperature of 37°C and an intermittent shaking protocol (alternating 10 sec of linear shaking with 1 mm amplitude and 50 sec without shaking) to support the growth of aerobic as well as microaerophilic strains.

#### Growth experiments with spent media

Spent media was obtained from fresh BHI+ inoculated with a 1:100 dilution from an OD600=0.3 suspension from each individual strain from the PNC8. These were incubated O/N at 37°C with orbital shaking at 250 rpm. After the incubation period, cultures were centrifuged at 4000 rpm and filtered using sterile filters (Whatman Puradisc 30, 0.2 micrometer pore size), to remove the cells. The pH was measured, and an aliquot of each spent medium was separated to adjust its pH back to 7.2-7.4 (Fresh BHI+ pH) by adding HCl 1M or NaOH 1N when necessary. The 3 types of spent media obtained were used to grow every single strain, alongside with fresh BHI+ as positive control. Experiments were conducted in 96-well plates, and OD600=0.3 suspensions were diluted 1:30 in the obtained media. 100 µL of mineral oil were added on top in order to avoid both evaporation and condensation. As described above, bacterial growth (as determined by OD600) was monitored every 10 min for around 24h with an automated plate reader (Tecan Nano M+) using an incubation temperature of 37°C and an intermittent shaking protocol.

#### Pairwise co-cultivation experiments

Pairwise cocultures were also conducted in a 96-well plate format. Again, after making PBS suspensions at OD600=0.3 for each strain, pairs of strains were inoculated 1:30 each in the media used (150 microL per well), 100 microL mineral oil was added, and bacterial growth was monitored as described in the previous sections.

#### Data processing

Microbial growth curves in 96-well plate format were analyzed using custom MATLAB scripts (Version R2021a) following procedures described previously ^63^. Briefly, raw OD600 time courses from each well were blank-corrected and smoothed using a moving average window with size 3. Area-under-the-growth-curve (AUC) was calculated using the *trapz* function, and the maximal growth rate of each curve during exponential growth was calculated using a slide window of six consecutive time points and an OD600 threshold of > 0.02.

### Exometabolome quantification

Exometabolome quantification was performed as follows: for each strain of interest, two replicate spent-media samples were generated as described in the spent media experiments above (using 7 mL of culture per cultivation tube) and stored at −20°C until further usage. Uninoculated media were processed in parallel to serve as a fresh-media reference. Relative metabolite concentrations in these supernatants (relative to fresh media) were subsequently quantified using gas-chromatography coupled to mass spectrometry (GC-MS) as described previously ^64^. Briefly, supernatant samples were extracted and derivatized following established protocols and measured using an Agilent 7890A GC coupled to a 5975C mass detector ^64^. Metabolites were identified manually with the TagFinder software using the reference library mass spectra and retention indices housed in the Golm Metabolome Database ^65^. In accordance with reporting recommendations ^66^, the parameters used for the peak annotation of all detected metabolites can be found in data set 3 (see below). In addition, L-lactate concentrations (relative to fresh media) were quantified using a commercial enzymatic assay (K-DLATE, Megazyme) following the manufacturer’s instructions. Subsequent data analysis was performed using custom MATLAB scripts (Version R2021a).

## Data availability

The PNC8 strain genome sequences generated in this study were deposited in the NCBI database (BioProject ID: PRJNA1175893). The data sets generated in this study are provided in six individual files as described below:

- data set 1 – KEGG module information for PNC8 strains and core nasal microbiome
- data set 2 – growth capacity of individual PNC8 strains across 23 *in vitro* conditions
- data set 3 – exometabolomics data of PNC8 members and additional strains
- data set 4 – spent-media experiments
- data set 5 – co-cultivation simulations
- data set 6 – pairwise co-cultivation experiments

## Supporting information

Supplementary text and figures are provided as a single document containing:

- **Supplementary text 1: Rational design of a defined consortium that recapitulates the *in vivo* composition of the pig nasal microbiota**
- **Supplementary text 2: Simulating competitive and non-competitive pairwise co-cultures**
**- Supplementary figures 1-13**

## Supporting information

data set 1

data set 2

data set 3

data set 4

data set 5

data set 6

## Acknowledgements

We thank Marcelo Gottschalk for kindly providing the bacterial strains *S.suis* P1/7 and *A.pleuropneumoniae* 4074.

## Author contributions

Conceived and designed the study: LBL, VA, KK. Performed experiments: LBL, OR, IFS. Performed analyses: LBL, OR, POG, IFS, KK. Supervised analyses: FCF, MS, VA, KK. Wrote manuscript with contributions from all authors: LBL, OR, KK.

