## Supplementary Text for "*In vitro* metabolic interaction network of a rationally designed nasal microbiota community"

*****: shared-first authors

**#correponding authors**:

**Supplementary text**

**Supplementary text 1 – Rational design of a defined consortium that recapitulates the *in vivo* composition of the pig nasal microbiota**

To develop a defined nasal microbiota consortium that recapitulates the *in vivo* composition of the pig nasal microbiota, we aimed to identify those taxa that are highly abundant in the nasal microbiota of healthy piglets. To obtain a broad picture of the nasal microbiota in healthy piglets, we analyzed 16S sequencing nasal swabs from 94 animals sampled at various farms from previous studies in our laboratory (processed jointly as described here ^1^). Since we were interested in nasal colonizers that can be grown under aerobic conditions, we excluded taxa belonging to *Clostridiales*/*Bacteroidales* (gut-associated anaerobic microbes which can frequently be found in nasal microbiota samples ^1^) from further analyses and re-calculated the relative abundance of all other taxa accordingly.

First, we analyzed the compositional complexity of these nasal microbiota samples in terms of the number of abundant taxa. We found that individual samples were typically dominated by a small set of highly abundant taxa at genus level (**Supplementary Figure 1A**). This low number of genera dominating the nasal microbiota of individual animals is in line with recent observations from human nasal microbiota samples ^2,3^ .

It is conceivable that although few taxa make up the bulk of individual nasal microbiota samples, the identity of these taxa may still vary widely across individuals. To assess this question, we next determined the relative abundance of different taxa (at genus level) across individuals (**Supplementary Figure 1B**). Consistent with previous reports ^4^, we found that *Moraxella*, *Lactobacillus*, and *Streptococcus* were the most abundant genera with close to 100% prevalence. In addition, we detected five other genera (*Rothia, Bergeyella, Glaesserella, Staphylococcus, Neisseria*), which were ranked just below (rank 4-8). Beyond these eight genera, we largely detected genera associated with the gut microbiota (e.g. *Trepanoma*, *Escherichia*), genera with ambiguous annotation, or genera with low relative abundance/prevalence. As a compromise between completeness (i.e. how many of the genera found in the pig nasal microbiota are included in the consortium) and size (i.e. enabling *in vitro* tractable experiments in 96-well plate format), we decided to consider the highest ranked eight genera for the defined consortium. Collectively, these eight genera represent the majority of the pig nasal microbiota (in terms of relative abundance) in individual animals (median: 64% summed relative abundance, with many samples reaching > 90% summed relative abundance, see **Figure 1C** in main text) both in this data set, as well as in five additional publicly available data sets of pig nasal microbiota samples (**Supplementary Figure 1C**). Interestingly, there is a high overlap between the genera selected here and the most abundant taxa in the human nasal microbiota. For example, *Moraxella*, *Staphylococcus*, *Neisseria*, and *Streptococcus*, were also identified among the 10 most abundant genera in the human nasal microbiota ^3^ (and all eight genera selected here were among the top 40 most abundant genera).

Next, we aimed to assemble a defined consortium that covers all eight genera selected above. Here, we decided to select one strain per genus, making use of our internal collection of strains that we had previously isolated from the nasal cavity of healthy piglets. Wherever possible, we prioritized strains with prior information regarding virulence traits, availability of a fully sequenced genome, and the ability to grow *in vitro*. For example, to represent the *Moraxella* and *Glaesserella* genera we selected *Moraxella pluranimalium* strain LG6-2 and *Glaesserella parasuis* strain F9, which have high serum susceptibility and low phagocytosis resistance ^5,6^. The full list of selected strains with additional information is summarized in **Supplementary Table 1**.

**Supplementary text 2** **–** **Simulating competitive and non-competitive pairwise co-cultures**

Here, we aimed to examine the impact of competition for nutrients on the dynamics of microbial pairwise co-cultures in more detail. Towards this end, we performed computational simulations using two different models (**Supplementary Figure 9A**).

Model 1 (**no-competition model**) is a co-culture of two strains, S_1_ and S_2_, which each have a different growth-limiting metabolite (M_1_ and M_2_, respectively). Using Monod kinetics ^7^ (and omitting a maintenance term to simplify the model), this model can be described by four ordinary differential equations (ODEs):

$$\frac{S_{1}}{dt}=\mu_{max,1}\cdot\frac{M_{1}}{K_{S1,M1}+M_{1}}\cdot S_{1}$$

**(eq 1.1)**

$$\frac{S_{2}}{dt}=\mu_{max,2}\cdot\frac{M_{2}}{K_{S2,M2}+M_{2}}\cdot S_{2}$$

**(eq 1.2)**

$$\frac{M_{1}}{dt}={-\mu}_{max,1}\cdot\frac{M_{1}}{K_{S1,M1}+M_{1}}\cdot S_{1}\cdot\frac{1}{Y_{S1,M1}}$$

**(eq 1.3)**

$$\frac{M_{2}}{dt}={-\mu}_{max,2}\cdot\frac{M_{2}}{K_{S2,M2}+M_{2}}\cdot S_{2}\cdot\frac{1}{Y_{S2,M2}}$$

**(eq 1.4)**

Where S1 and S2 denote the biomass of strain 1 and 2, μ_max,1_ and μ_max,2_ denote their respective maximal growth rates, M_1_ and M_2_ denote the concentration of the growth-limiting metabolites, K_S1,M1_ and K_S2,M2_ denote the respective Monod constants, and Y_S1,M1_ and Y_S2,M2_ denote the biomass yields for each strain-metabolite combination.

Model 2 (**competition model**) is a co-culture of two strains, S_1_ and S_2_, which both compete for the same growth-limiting metabolite M_1_. This model can be described by three ODEs:

$$\frac{S_{1}}{dt}=\mu_{max,1}\cdot\frac{M_{1}}{K_{S1,M1}+M_{1}}\cdot S_{1}$$

**(eq 2.1)**

$$\frac{S_{2}}{dt}=\mu_{max,2}\cdot\frac{M_{1}}{K_{S2,M1}+M_{1}}\cdot S_{2}$$

**(eq 2.2)**

$$\frac{M_{1}}{dt}={-\mu}_{max,1}\cdot\frac{M_{1}}{K_{S1,M1}+M_{1}}\cdot S_{1}\cdot\frac{1}{Y_{S1,M1}} -\mu_{max,2}\cdot\frac{M_{1}}{K_{S2,M1}+M_{1}}\cdot S_{2}\cdot\frac{1}{Y_{S2,M1}}$$

**(eq 2.3)**

Where K_S1,M1_ and K_S2,M1_ denote the respective Monod constants for M_1_, and Y_S1,M1_ and Y_S2,M1_ denote the respective biomass yields for each strain.

To simulate the behavior of these models, we used parameter ranges that reflected the experimentally observed parameter ranges in this study, namely μ_max_ between 0.1 and 1.7h^-1^, biomass yields between 0.1 and 1 g/g, and used a simulation duration of 24h (which matches the approximate duration of the experimental *in vitro* cultivations performed in this study). Initial biomass and Monod constants were set to 0.01 g/L, and initial metabolite concentrations were set to 1 g/L.

We performed simulations with 1000 strain pairs whose kinetic parameters were uniformly sampled from the aforementioned parameter ranges. For each strain pair, we simulated co-cultivation without or with competition using the models described above (adding equal amounts of starting biomass, namely 0.01 g/L), as well as cultivation of each strain individually (**Supplementary Figure 9B)**. As expected, without metabolite competition the measured maximal co-culture biomass matches exactly the predicted co-culture biomass (i.e. predicted by summing the maximal biomass of each individual strain culture). In contrast, co-cultures with competition invariably show lower-than-expected biomass production (i.e. co-culture biomass production is lower than the summed maximal biomass of each individual strain culture). These findings make intuitive sense – if two strains need to compete for the same metabolite, their summed biomass naturally will be lower than if each strain is grown separately using the same initial metabolite concentration – and also match experimental observations from previous studies ^8^.

The simulated pairwise co-cultures described here provided a baseline model of the expected co-culture dynamics for strains that compete for a single growth-limiting metabolite (in absence of any additional interaction mechanisms). Next, we wanted to test whether these simulations (in which strains with wildly different growth rates and yields were pitted against each other) could also reveal which strain outcompetes the other based on their cultivation time courses alone (as postulated e.g. here ^9^, despite the inherent limitations of such an approach ^10^). Specifically, we hypothesized that when co-cultivating two competing strains, the most obvious way by which one strain can outcompete the other is by growing much faster. Moreover, we speculated that the more one strain outcompetes the other, the more the co-culture dynamics will recapitulate its individual culture. To test these conjectures, we calculated for each simulated strain pair the final relative abundance of each strain, as well as the relative distance between co-culture and single culture time courses. These analyses revealed indeed a strong correlation between the difference in μ_max_ and not only the final relative abundance of each strain across simulated strain pairs (**Supplementary Figure 9C**), but also the relative distance of their individual cultures to the co-culture dynamics (**Supplementary Figure 9D**). This strong correlation was absent in non-competing strain pairs (gray circles in the same plots).

Taken together, these simulations provide two key predictions for the expected behavior of co-cultures of strain pairs that compete for growth-limiting metabolites. First, for any given strain pair, the measured biomass production is predicted to be substantially below the summed biomass production of each strain grown in isolation. Second, in absence of additional effects, the faster-growing strain will invariably outcompete the slower one, and the larger the growth rate difference, the more the resulting co-culture will recapitulate the growth curve of the faster growing strain.

**Supplementary Tables**

**Supplementary Table 1** **–** **Strains used in this study**

| **Genus** | **Strain** | **Source**  **(References at the end of the SI)** | **PNC8 strain** |
| --- | --- | --- | --- |
| *Moraxella* | *Moraxella pluranimalium* LG6-2 | ^5^ | **Yes** |
| *Lactobacillus* | *Lactobacillus odoratitofui* KD9-5 | This study | **Yes** |
| *Streptococcus* | *Streptococcus pluranimalium* LG3-6 | ^11^ | **Yes** |
| *Rothia* | *Rothia nasimurium* UK1-9 | This study | **Yes** |
| *Bergeyella* | *Bergeyella zoohelcum* AR-9 | ^12^ | **Yes** |
| *Glaesserella* | *Glaesserella parasuis* F9 | ^13^ | **Yes** |
| *Staphylococcus* | *Staphylococcus aureus* EJ41-2 | This study | **Yes** |
| *Neisseria* | *Neisseria shayeganii* GM3-3 | This study | **Yes** |
| *Staphylococcus* | *Staphylococcus succinus* GW5-1 | This study | **No** |
| *Rothia* | *Rothia nasimurium* VL1-3 | This study | **No** |
| *Streptococcus* | *Streptococcus pluranimalium* GW6-1 | This study | **No** |
| *Glaesserella* | *Glaesserella parasuis* Nagasaki | ^14^ | **No** |
| *Streptococcus* | *Streptococcus suis* P1/7 | ^15^ | **No** |
| *Actinobacillus* | *Actinobacillus pleuropneumoniae* 4074 | ^16^ | **No** |

**Supplementary Table 2 –** **Cultivation media used in this study**

| **Medium name** | **Composition** |
| --- | --- |
| BHI | Brain Heart Infusion broth (53286, Sigma-Aldrich) |
| BHI+ | Brain Heart Infusion broth + 80 µg/mL NAD^+^ (N0632, Sigma-Aldrich) + 1% pig serum heat-deactivated (P9783, Sigma-Aldrich) |
| M9 base salts | 11.33 g/L sodium phosphate dibasic heptahydrate (S9390, Sigma-Aldrich) + 3 g/L potassium phosphate monobasic (P0662, Sigma-Aldrich) + 0.5 g/L sodium chloride (1.06404.1000, Millipore) + 1 g/L ammonium chloride (12125-02-9, Millipore) |
| M9 + Glc | M9 base salts + 2 g/L glucose (16301, Sigma-Aldrich) |
| M9 + Glc + Vit | M9 base salts + 2 g/L glucose + 2% RPMI vitamin solution (R7256, Sigma-Aldrich) + 80 µg/mL NAD^+^ + 1 mg/mL pyridoxal-HCl (P9130, Sigma-Aldrich) + 1 mg/mL pyridoxamine-2HCl (P9158, Sigma-Aldrich) |
| M9 + Glc + CAA | M9 base salts + 2 g/L glucose + 2 g/L Casamino acids (N4642, Sigma-Aldrich) |
| M9 + Glc + RPMI | M9 base salts + 2 g/L glucose + 1% RPMI amino acid solution (R7131, Sigma-Aldrich) + 2 mM L-Glutamine (G7513, Sigma-Aldrich) |
| M9 + Glc + CAA + Vit | M9 base salts + 2 g/L glucose + 2 g/L CAA + 2% RPMI vitamin solution + 80 µg/mL NAD^+^ + 1 mg/mL pyridoxal-HCl + 1 mg/mL pyridoxamine-2HCl |
| M9 + Glc + RPMI + Vit | M9 base salts + 2 g/L glucose + 1% RPMI amino acid solution + 1% L-Glutamine + 2% RPMI vitamin solution + 80 µg/mL NAD^+^ + 1 mg/mL pyridoxal-HCl + 1 mg/mL pyridoxamine-2HCl |
| M9 + Glc + CAA + Vit + pig serum | M9 base salts + 2 g/L glucose + 2 g/L CAA + 2% RPMI vitamin solution + 80 µg/mL NAD^+^ + 1 mg/mL pyridoxal-HCl + 1 mg/mL pyridoxamine-2HCl + 1% pig serum heat-inactivated |
| M9 + Glc + CAA + Vit + yeast extract | M9 base salts + 2 g/L glucose + 2 g/L CAA + 2% RPMI vitamin solution + 80 µg/mL NAD^+^ + 1 mg/mL pyridoxal-HCl + 1 mg/mL pyridoxamine-2HCl + 0.2 g/L yeast extract (70161, Sigma-Aldrich) |
| M9 + Glc + CAA + Vit + BHI | M9 base salts + 2 g/L glucose + 2 g/L CAA + 2% RPMI vitamin solution + 80 µg/mL NAD^+^ + 1 mg/mL pyridoxal-HCl + 1 mg/mL pyridoxamine-2HCl + 1% BHI |
| M9 + Glc + CAA + Vit + THB | M9 base salts + 2 g/L glucose + 2 g/L CAA + 2% RPMI vitamin solution + 80 µg/mL NAD^+^ + 1 mg/mL pyridoxal-HCl + 1 mg/mL pyridoxamine-2HCl + 1% THB |
| M9 + Glc + CAA + Vit + MRS | M9 base salts + 2 g/L glucose + 2 g/L CAA + 2% RPMI vitamin solution + 80 µg/mL NAD^+^ + 1 mg/mL pyridoxal-HCl + 1 mg/mL pyridoxamine-2HCl + 1% MRS |
| M9 + NAD^+^ + pig serum | M9 base salts + 80 µg/mL NAD^+^ + 1% pig serum heat-inactivated |
| MH | Müller-Hinton broth (212322, BD) |
| THY | Todd-Hewitt Broth (T1438, Millipore) + 0.2 g/L yeast extract |
| MRS | De Man, Rogosa and Sharpe broth (69966, Sigma-Aldrich) |
| LB | Luria-Bertani broth (L3022, Sigma-Aldrich) |
| HPLM | Human Plasma-Like Medium (A4899101, Gibco) |
| BHI + NAD^+^ + CAA | BHI + 80 µg/mL NAD^+^ + 2 g/L CAA |
| BHI + NAD^+^ + yeast extract | BHI + 80 µg/mL NAD^+^ + 0.2 g/L yeast extract |
| BHI + Vit | BHI + 1% RPMI vitamin solution + 80 µg/mL NAD^+^ + 1 mg/mL pyridoxal-HCl + 1 mg/mL pyridoxamine-2HCl |
| SNM30 (adapted from ^17^) | M9 base salts + 3.3 g/L glucose + amino acid solution (composed of 0.13 g/L alanine (A7627, Sigma-Aldrich) + 0.21 g/L arginine (A5131, Sigma-Aldrich) + 0.015 g/L cysteine (C1726, Sigma-Aldrich) + 0.19 g/L glutamate (49621, Sigma-Aldrich) + 0.11 g/L glycine (G7126, Sigma-Aldrich) + 0.1 g/L histidine (H8125, Sigma-Aldrich) + 0.4 g/L leucine (L8000, Sigma-Aldrich) + 0.27 g/L lysine (L5626, Sigma-Aldrich) + 0.17 g/L ornithine (O2375, Sigma-Aldrich) + 0.25 g/L phenylalanine (78019, Sigma-Aldrich) + 0.17 g/L proline (P0380, Sigma-Aldrich) + 0.13 g/L serine (S4500, Sigma-Aldrich) + 0.24 g/L threonine (T8625, Sigma-Aldrich) + 0.04 g/L tryptophan (T0254, Sigma-Aldrich) + 0.12 g/L valine (V0500, Sigma-Aldrich)) + organic acids solution (composed of 0.04 g/L citrate (C0759, Sigma-Aldrich) + 0.008 g/L fumarate (F1506, Sigma-Aldrich) + 0.014 g/L maleate (M5757, Sigma-Aldrich) + 0.11 g/L pyruvate (P2256, Sigma-Aldrich) + 0.54 g/L succinate (S2378, Sigma-Aldrich)) + 0.3 g/L urea (51546, Sigma-Aldrich) + 1% RPMI vitamin solution + 80 µg/mL NAD^+^ + 1 mg/mL pyridoxal-HCl + 1 mg/mL pyridoxamine-2HCl |
| SNM30 + Mucin | 75% v/v SNM30 (see above) + 25% v/v of 20 g/L porcine stomach mucin stock solution (M2378, Sigma-Aldrich, final concentration 5 g/L, stock solution prepared following previously established protocols ^18^) |

**Supplementary Figures**

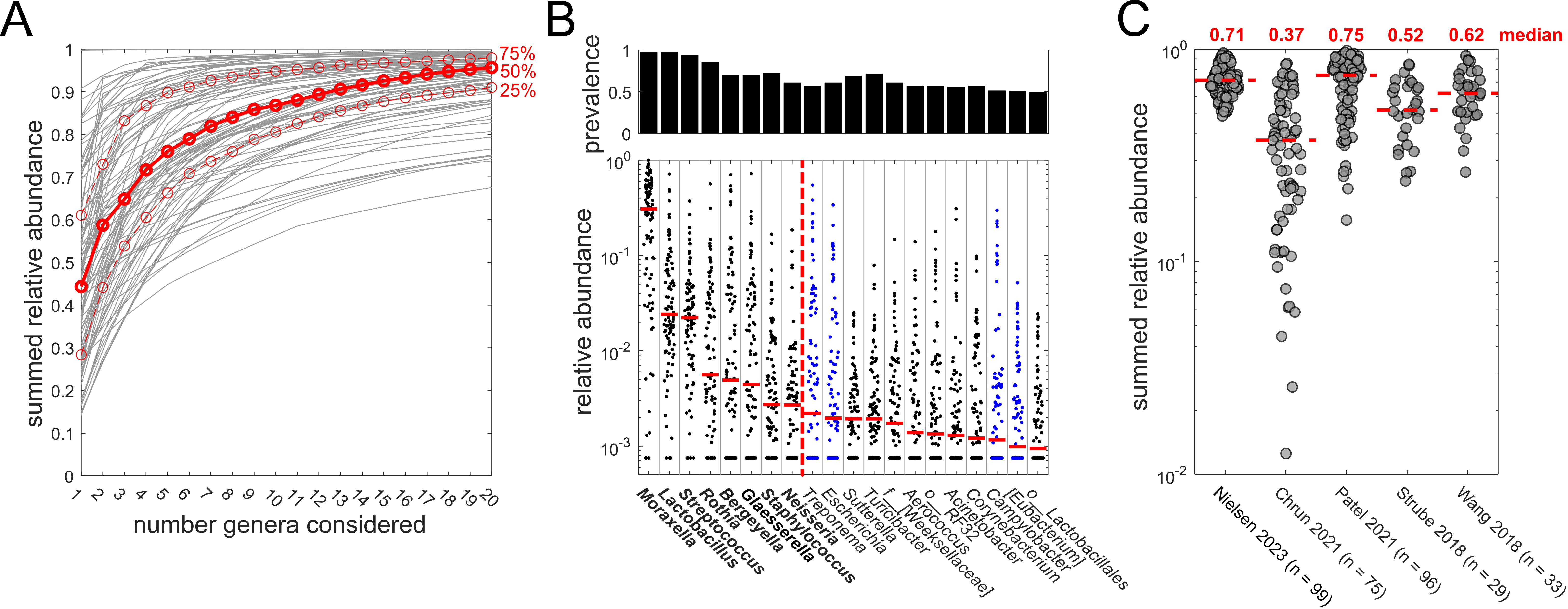

**Supplementary Figure 1. Rational design of the Porcine Nasal Consortium (PNC8) using *in vivo* nasal microbiota samples.** **A)** Summed relative abundance as a function of the number of taxa considered at genus level in nasal microbiota samples obtained from 94 piglets (after removing gut-microbiota associated taxa, see **Supplementary Text 1** and ^1^ for details). Gray lines denote samples from individual animals. Red dashed lines denote the 75 and 25 percentiles, and the red continuous line denotes the median. **B)** Prevalence (fraction of samples with relative abundance > 0.1%, top) and relative abundance (bottom) of the twenty most abundant genera (sorted by median abundance across samples, shown as red horizontal lines). Each circle denotes an individual animal. Blue: respective genus is prevalent (>10% prevalence, as described here ^1^) in a reference pig gut microbiota data set ^19^. Vertical red dashed line: cut-off of genera included in the PNC8 (shown in bold). Wherever the taxonomical assignment was not possible at genus level, the lowest resolved annotation is shown. **C)** Summed relative abundance of PNC8 genera in five publicly available pig nasal microbiota data sets ^20–24^ (using in each case the control group animals, n denotes the number of samples used for each data set). Red dashed lines: median summed relative abundance (values are listed above).

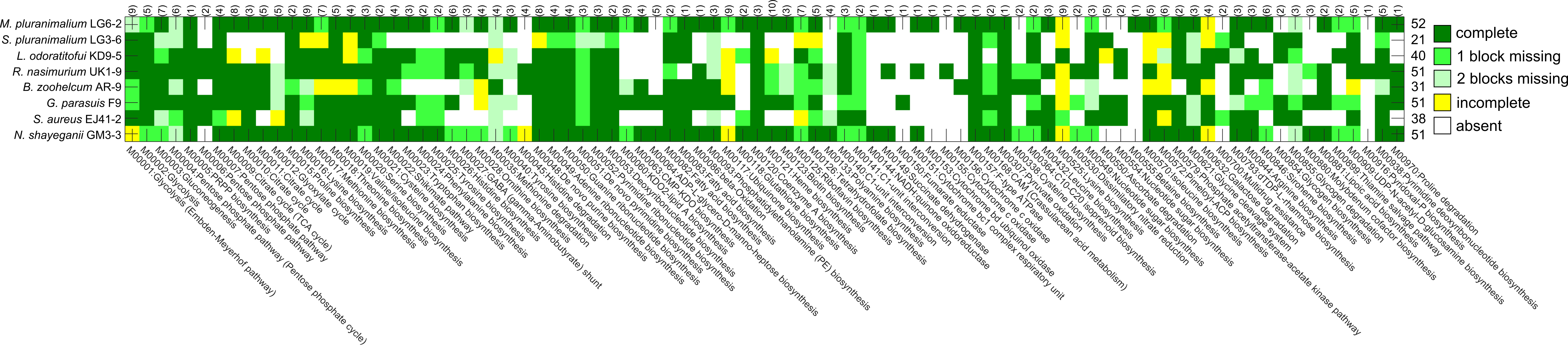

**Supplementary Figure 2.** **KEGG module completeness in the PNC8 strains.** Only modules that were complete in at least one strain were considered. Vertical numbers in brackets: number of blocks within respective module. Horizontal numbers: number of complete KEGG modules in each strain. KEGG module completeness was determined with the “reconstruct” function of the KEGG-mapper tool (available online at <https://www.genome.jp/kegg/mapper/>) as described in the methods section of the main text.

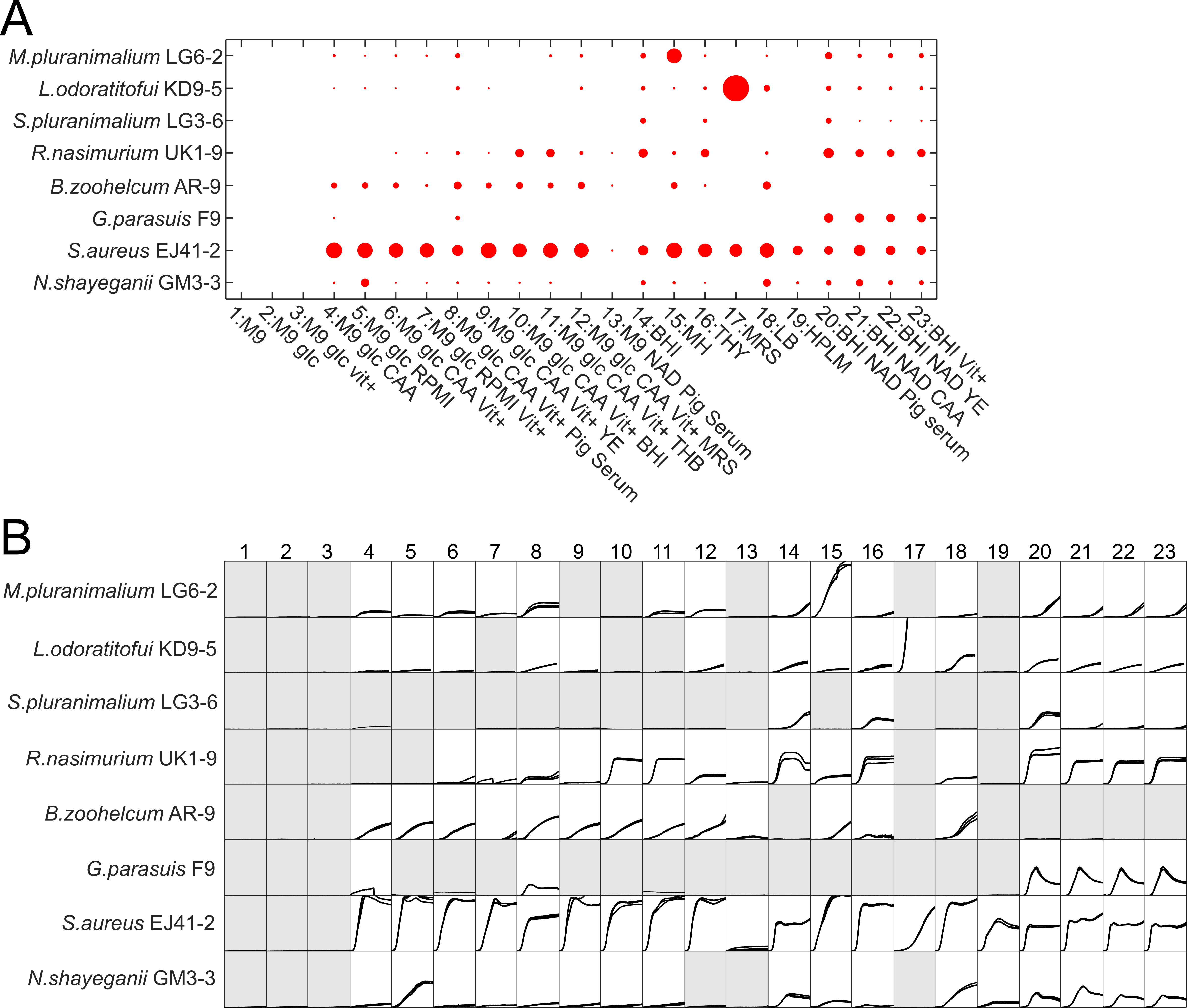

**Supplementary Figure 3**. **Growth patterns of PNC8 members across various *in vitro* conditions. A)** Maximal OD600 value (mean across 2-3 replicates) shown as red circle size for all eight PNC8 members in 23 different cultivation media. **B)** Growth curves of all PNC8 members in 23 different cultivation media (see **Supplementary Table 2** for list of used media). Each line denotes a replicate culture. Gray: no growth detected (maximal OD600 < 0.02). X-axis range (h): 0-24h. Y-axis range (OD600): 0 to 0.6.

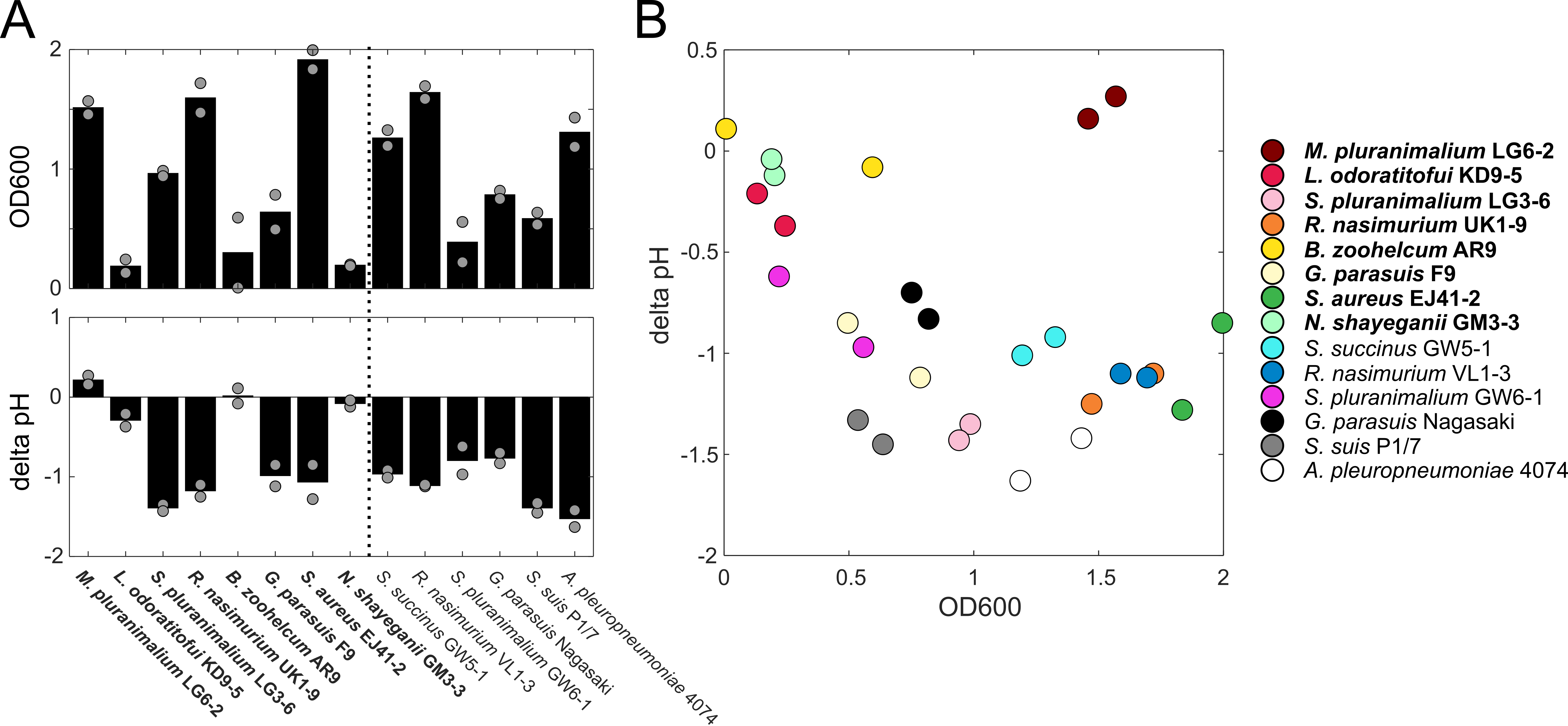

**Supplementary Figure 4. OD600 and pH changes of nasal microbiota members in BHI Suppl. media.** **A)** Final OD600 (top) and pH change compared to fresh media (bottom) of PNC8 strains (bold strain names) and additional nasal microbiota strains. Two replicate cultures were measured for each strain (shown as grey circles). Bars denote the respective mean. **B)** Final OD600 plotted against pH change for all replicate cultures.

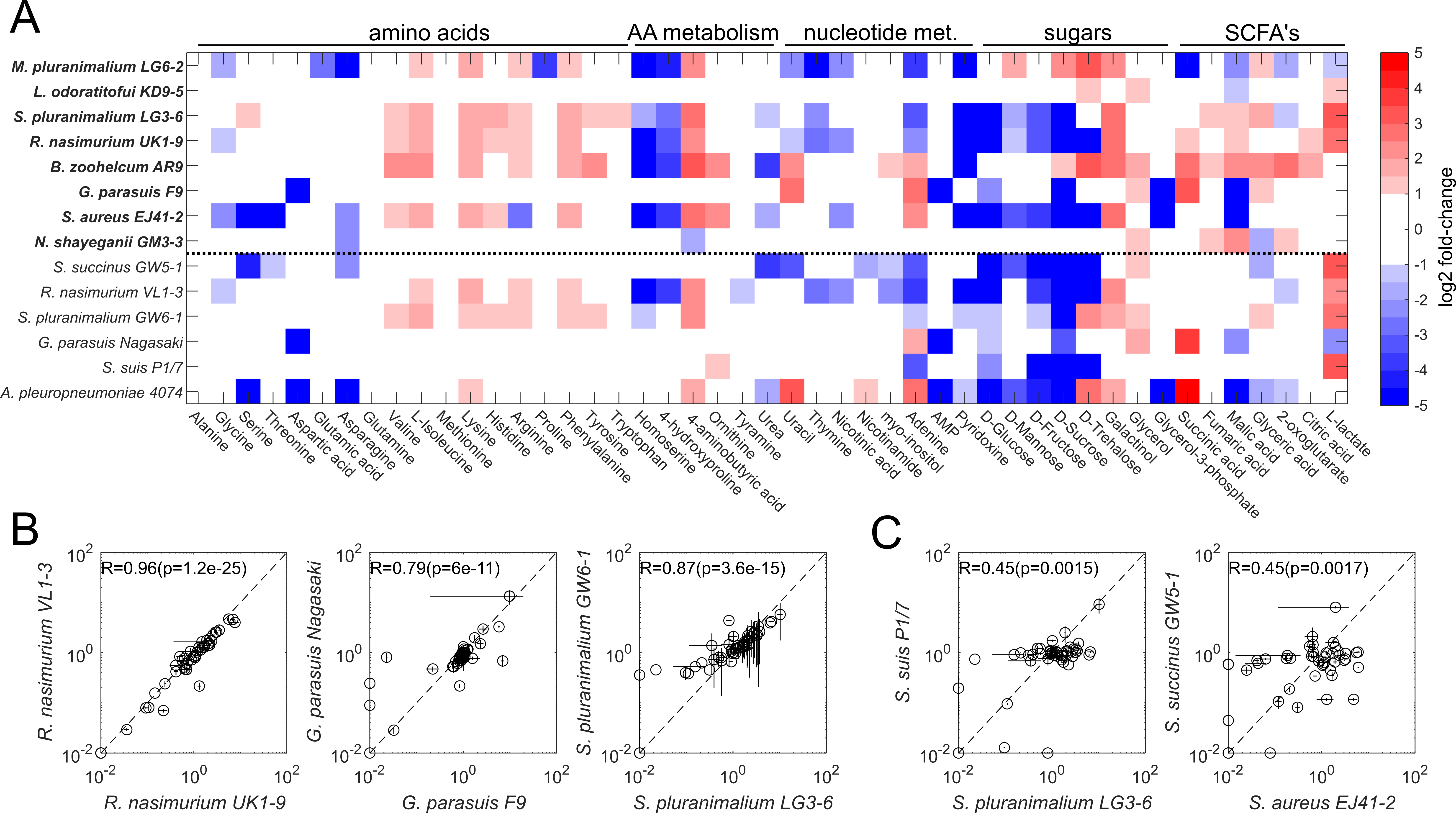

**Supplementary Figure 5. Exometabolome patterns of nasal microbiota members in spent media. A)** Heatmap of extracellular metabolite fold-changes (relative to the concentration in fresh BHI+ media) across all tested strains. Data show the average of two replicate cultures. Metabolites were sorted according to metabolite types. Each row denotes a nasal microbiota strain. Top 8 rows (in bold): PNC8 strains. Rows below dashed line: other nasal microbiota strains. **B)** Comparison of exometabolome patterns of strains belonging to the same species. Error bars denote standard deviation (n = 2). R denotes Spearman correlation (and p the corresponding p-value). **C)** Comparison of exometabolome patterns of strains belonging to the same genus.

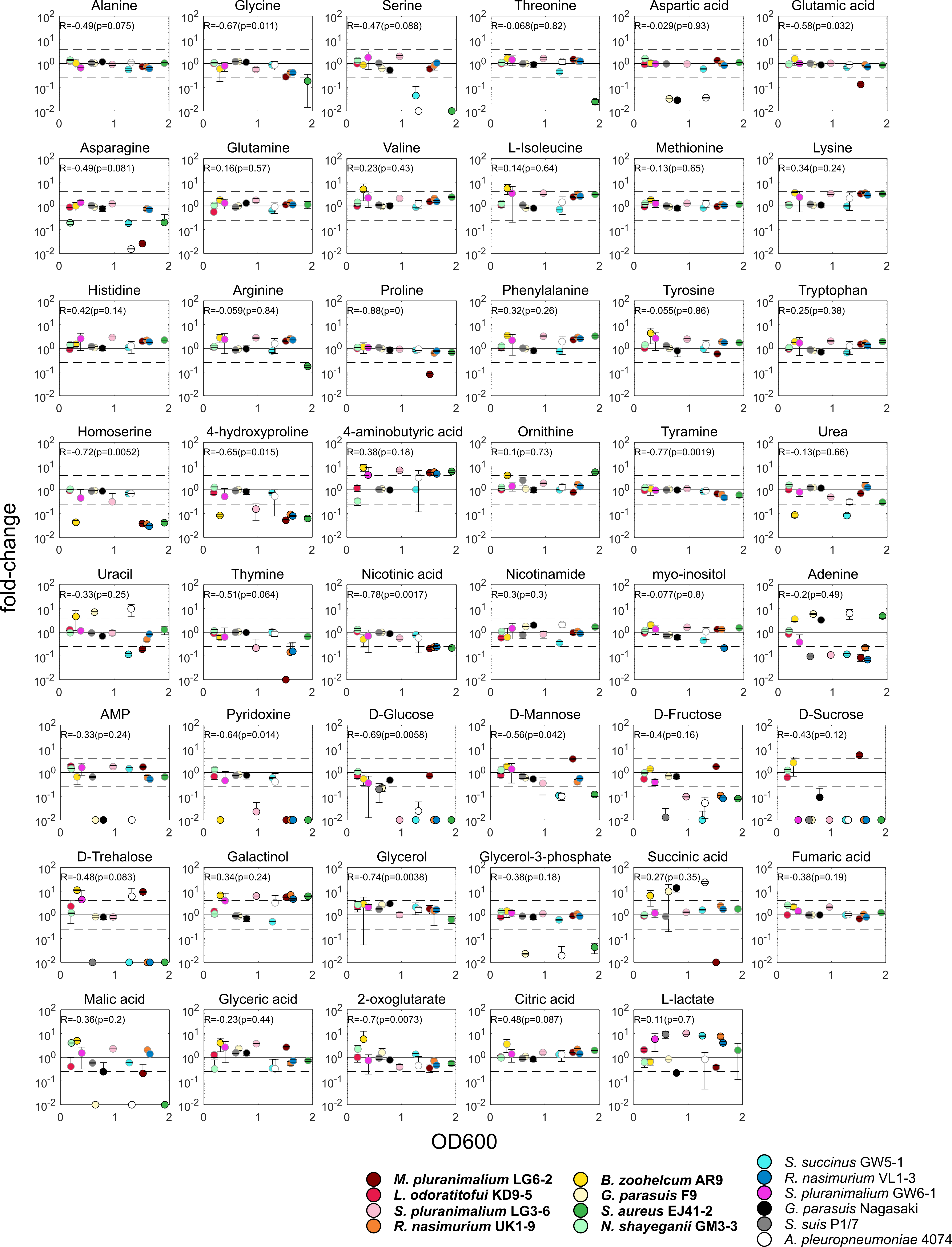

**Supplementary Figure 6. Exometabolite fold-change (compared to fresh media) plotted against final OD600 across all 14 tested strains.** Error bars denote standard deviation (n = 2). Horizontal dashed lines: four-fold change. R: Spearman correlation, p: corresponding p-value.

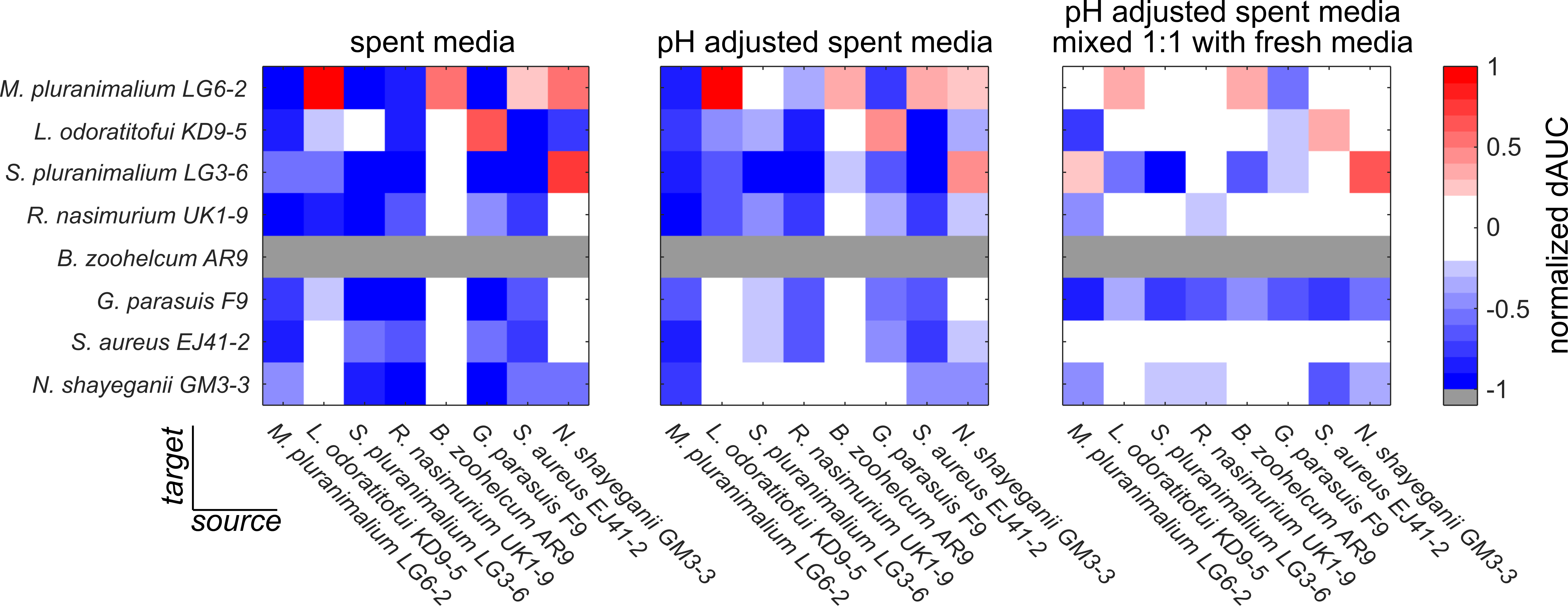

**Supplementary Figure 7. Normalized difference in area-under-the-growth-curves (dAUC). Normalized dAUC = (AUC_spent_ – AUC_fresh_) / AUC_fresh_) for all PNC8 strains**. Negative normalized dAUC values denote cases where a strain grows more poorly in a spent media compared to fresh media. Data show mean of 2-3 replicate cultures. Left: spent media. Middle: pH adjusted spent media. Right: pH adjusted spent media mixed 1:1 with 2x concentrated fresh BHI media (but no additional NAD+ and pig serum), where in most cases the normalized dAUC moves close to zero (indicating that growth in these cases is comparable to fresh media). One exception is *G. parasuis* F9, which grows poorly in all spent media likely due to technical reasons (this strain requires NAD+ supplementation for growth, which was omitted in the 2x concentrated fresh BHI media).

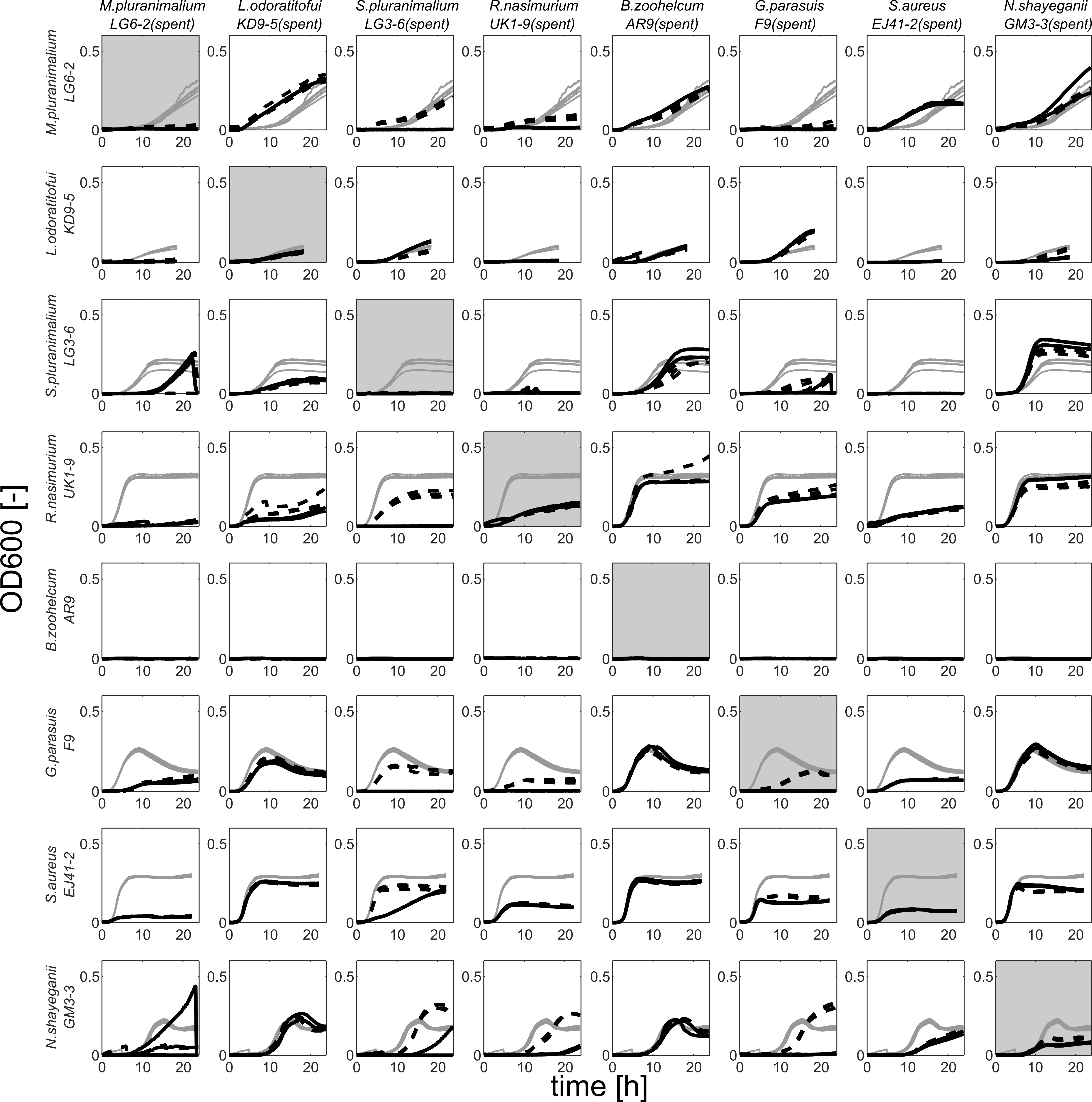

**Supplementary Figure 8**. **Spent-media growth curves.** Each column denotes spent media from each designated PNC8 strain, each column denotes the growth curve of each designated PNC8 strain in fresh media (gray lines), spent media (black continuous lines), and pH adjusted spent media (black dashed lines). Each curve denotes a replicate well (6 replicates for fresh media, 2-3 replicates for spent media experiments).

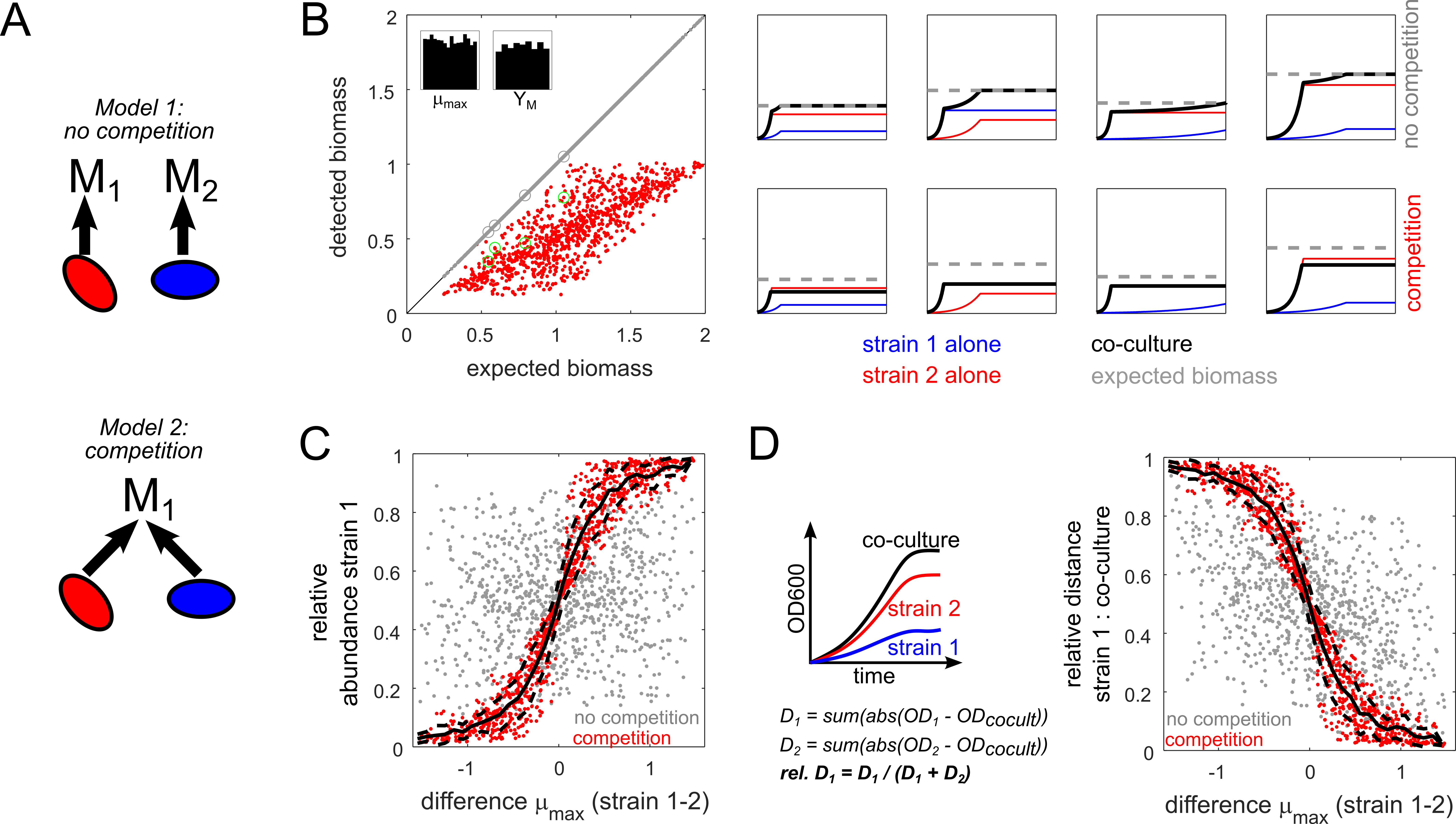

**Supplementary Figure 9. Simulations of competitive and non-competitive co-cultures. A)** Schematic of models. **B)** Left: Expected (i.e. sum of individual strains) versus detected biomass in 1000 non-competitive (gray circles, all laid out exactly on the diagonal) and competitive (red circles) strain pairs. Insets: distribution of parameter values for μ_max_ and biomass yield Y_M_. Right: example time courses for 4 randomly selected strain pairs (highlighted left with larger gray and green circles). Expected biomass was calculated as the sum of maximal biomass values for each individual strain. **C)** Difference in μ_max_ between strain 1 and 2 plotted against the relative abundance of strain 1 (at final time point). Black continuous line: moving average (window size 0.1 h^-1^). Dashed lines: respective standard deviation. **D)** Difference in μ_max_ between strain 1 and 2 plotted against the relative distance between the growth curves of strain 1 and the co-culture (calculated as indicated in the schematic). Black continuous line: moving average (window size 0.1 h^-1^). Dashed lines: respective standard deviation. All simulations were performed with MATLAB (Version R2021a) using the *ode45* solver (solver options: relative tolerance = 10^-8, absolute tolerance = 10^-10).

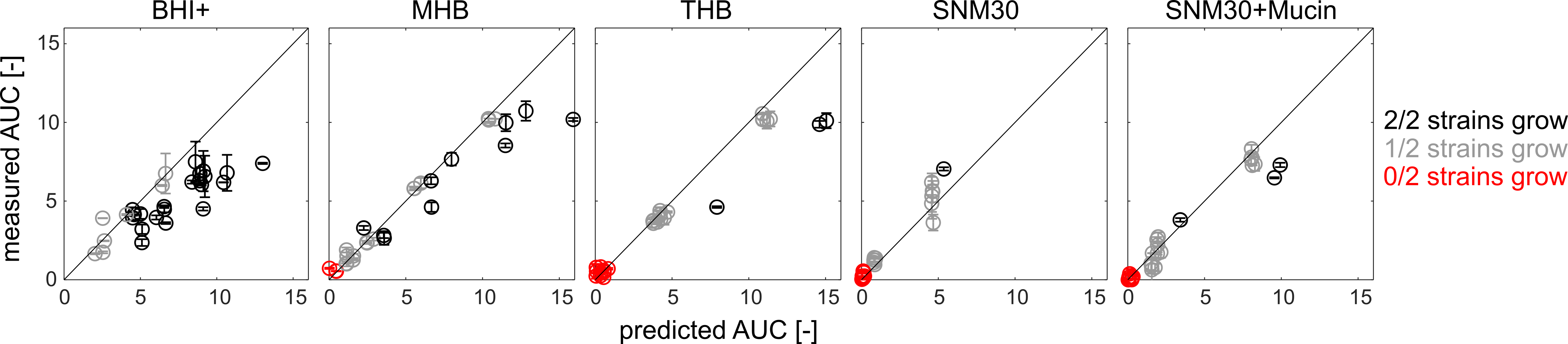

**Supplementary Figure 10. Predicted and measured biomass production across all tested conditions using an alternative biomass metric.** Predicted and measured Area-under-growth-curve (AUC) as an alternative biomass production metric for pairwise co-cultures of PNC8 strains across all tested conditions. Gray circles denote strain pairs in which only one of the two strains grows in isolation in the tested condition. Red circles denote strain pairs in which neither of the two strains grows in isolation in the tested condition. Error bars denote standard deviation (n = 2).

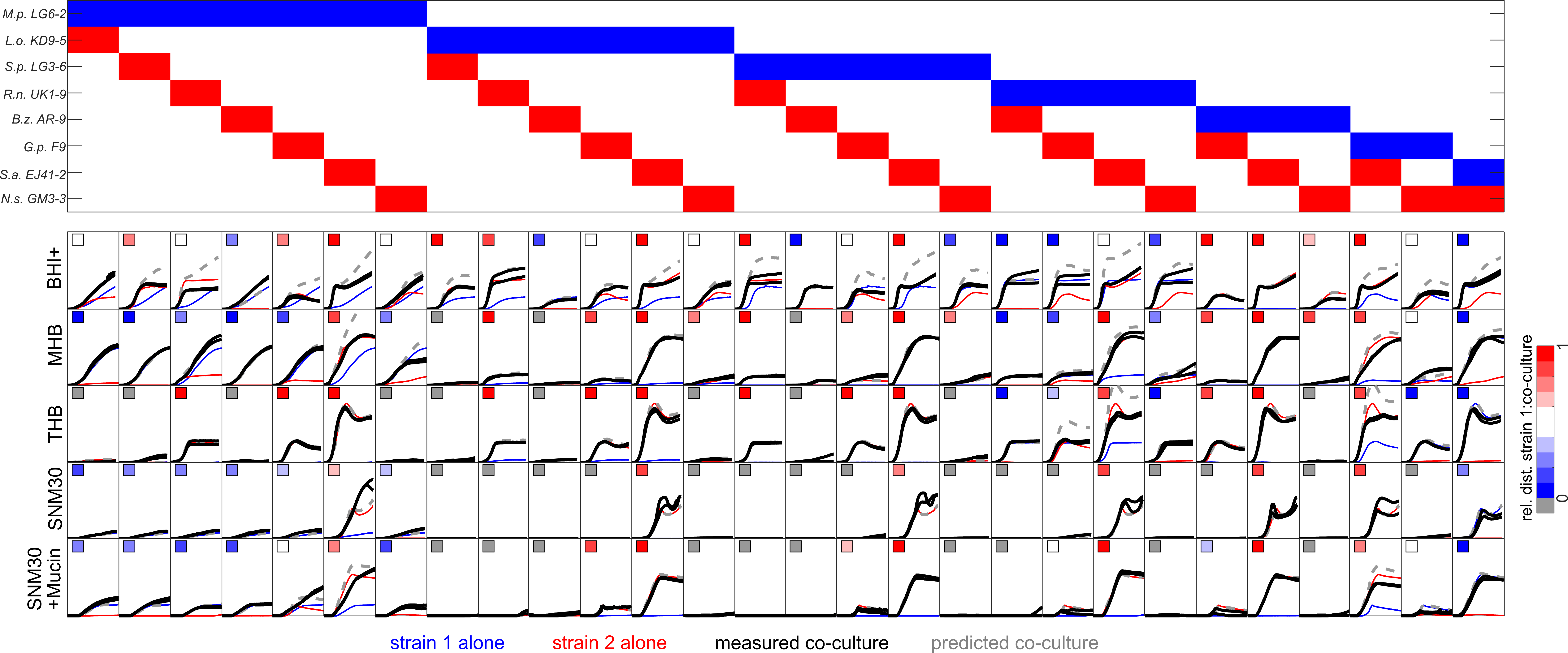

**Supplementary Figure 11. Co-cultivation growth curves for all PNC8 strain combinations.** Plots are sorted row-wise by strain combination and column-wise by condition. For any given plot, individual growth curves of strain 1 are shown in blue and strain 2 are shown in red (mean of two replicate curves). Black lines: co-culture. Gray dashed line: predicted co-culture (sum of individual strain time courses). Each black line denotes an individual replicate culture. Insets: relative distance of strain 1 to co-culture (calculated as described in **Supplementary Figure 9**). X-axis range (h): 0-24h. Y-axis range (OD600): 0 to 1.

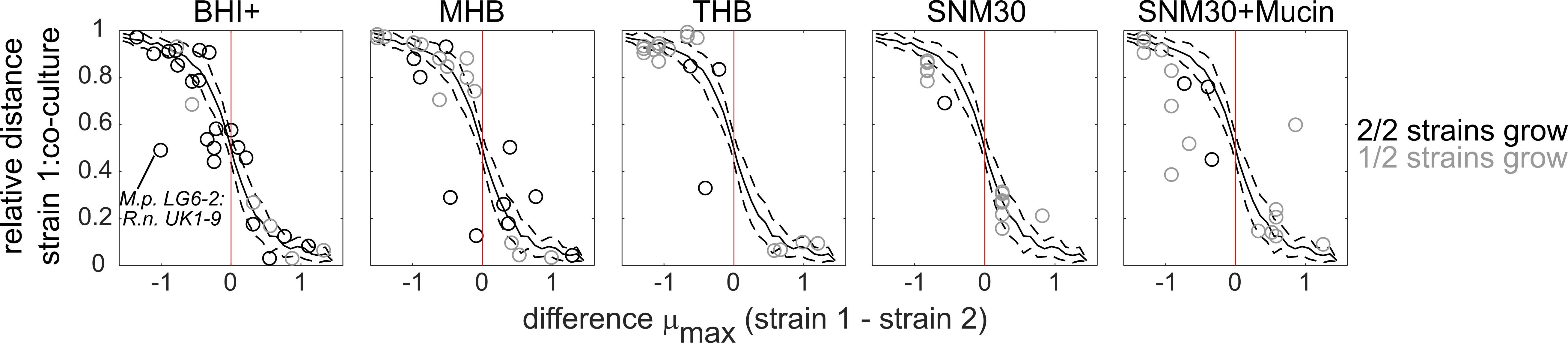

**Supplementary Figure 12.** Relative distance of strain 1 to co-culture (calculated as described in **Supplementary Figure 9**) plotted against the difference in maximal growth rate (μ_max_) between strain 1 and 2 (as determined when grown in isolation). Black circles: Both strains grow in isolation. Gray circles: only one of the two strains grows in isolation. Black lines: predicted relationship between growth rate difference and relative distance in 1000 simulated strain pairs competing for a single growth-limiting metabolite (see **Supplementary Text 2**). Moving average and corresponding standard deviation of these simulations are shown as continuous and dashed lines, respectively.

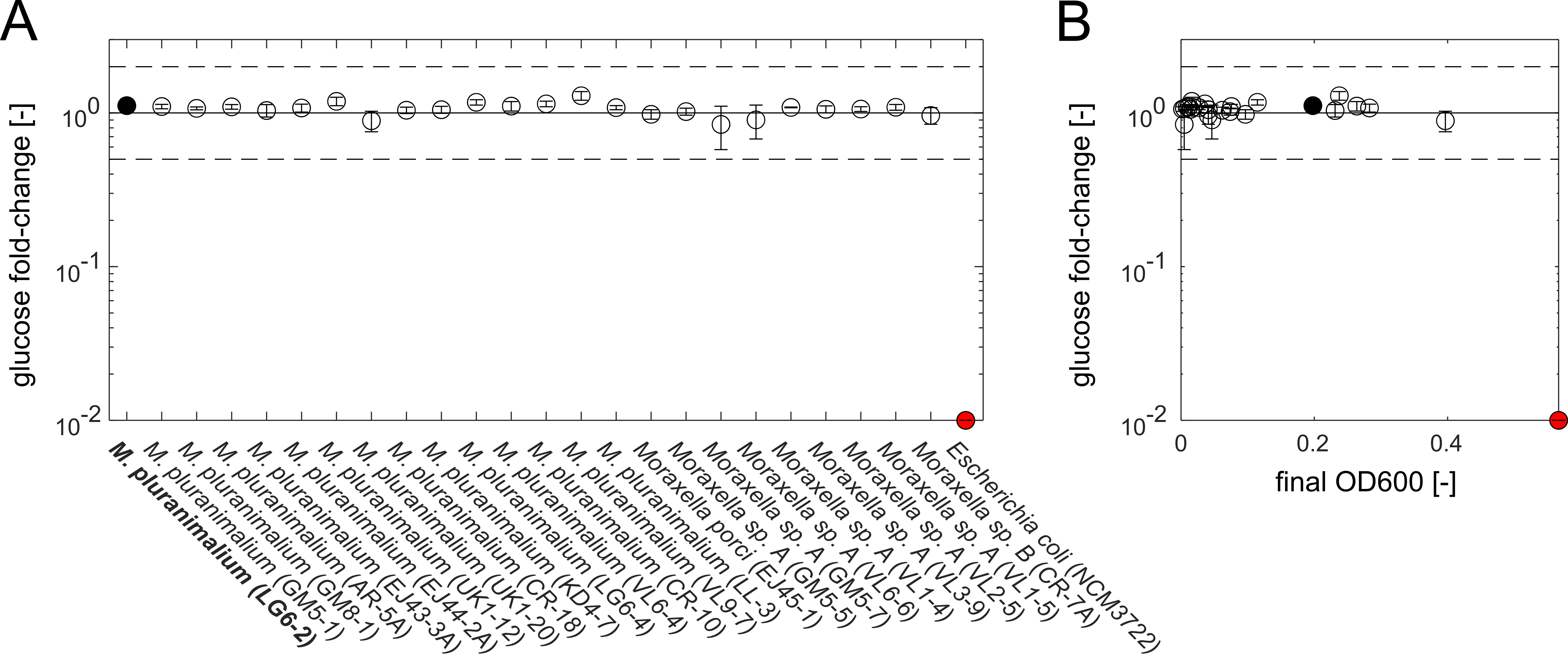

**Supplementary Figure 13. Quantifying glucose consumption of different *Moraxella* strains in BHI+. A)** Relative glucose concentration (relative to fresh BHI+) various *Moraxella* cultures (all strains were isolated from healthy piglets and originally described in ^5^) grown for 24h in BHI+ in 96-well plate format (using the same experimental protocol as in the methods section “*Growth experiments across metabolic environments*”). Relative glucose concentrations were determined with an enzymatic assay (GOPOD format, Cat K-GLUC, Megazyme) following the manufacturer’s instructions. Horizontal dashed lines: 2-fold change in glucose concentration. Black circle: strain included in the PNC8. Red circle: a laboratory *E.coli* strain serving as a positive control. Relative fold-changes below 0.01 were set to 0.01 to aid visualization. Error bars denote standard deviation (n = 3). **B**) Same data as in A), but plotted against final OD600 of each respective culture.
